# Cadherin Regulation of Endoplasmic Reticulum-Plasma Membrane Contact Sites

**DOI:** 10.1101/2025.10.15.682666

**Authors:** Sonam Lhamo, Navaneetha Krishnan Bharathan, William Giang, Kandice R. Levental, Cory L. Simpson, Stephanie Zimmer, Andrew P. Kowalczyk

## Abstract

The spatial organization and dynamics of the endoplasmic reticulum (ER) govern when and where ER tubules engage with other organelles and the plasma membrane. We previously found that ER tubules are closely associated with desmosomes, but the mechanisms of ER recruitment to these adhesive intercellular junctions were unclear. Here, we demonstrate that adherens junctions recruit ER tubules to intercellular junctions in a manner dependent upon E-cadherin association with α-catenin and vinculin. During cell-cell junction assembly, adherens junctions and ER appear nearly simultaneously at nascent cell-cell contacts, followed by desmosome formation. ER recruitment to cell-cell contacts allows the formation of ER-plasma membrane contact sites (ER-PMCS) and the assembly of a complex comprising adherens junctions, ER-PMCS, and desmosomes. Ablating adherens junctions disrupts this tripartite complex and perturbs global cellular lipid levels. Collectively, our findings identify cadherins as key organizers of ER-PMCS positioning and suggest that this complex integrates cellular mechanical elements with plasma membrane homeostasis.

## Introduction

The endoplasmic reticulum (ER) is a multifunctional organelle that regulates proteostasis, lipid homeostasis, and calcium signaling^1,2^. The ER performs many of these functions at specialized membrane contact sites where the ER is positioned within 10-25 nm of adjacent organelles or the plasma membrane to allow communication between cellular compartments^1–4^. ER morphology and spatial distribution influence its functions. For example, ER sheets in the perinuclear region are sites of protein synthesis and trafficking, whereas the peripheral tubular ER network forms contact sites with various organelles and the plasma membrane. ER-plasma membrane contact sites (ER-PMCS) influence cell migration, calcium signaling, and lipid metabolism^5–8^. However, the cellular mechanisms governing the assembly and spatial distribution of ER-PMCS are poorly understood.

Recent studies indicate that ER tubules associate with both cell-cell and cell-substrate adhesive interfaces^5,8,9^. We previously identified close physical and functional associations between the ER and the desmosome, an adhesive cell-cell junction^10,11^. Desmosomes are robust cell-cell adhesion complexes that are anchored to intermediate filaments and confer mechanical integrity to epithelial tissues and cardiac muscle^12–14^. Volume electron microscopy (EM) of epithelial cells revealed ER tubules in close proximity to desmosomes, often making physical contacts with both keratin filaments and the outer dense plaque of desmosomes^10^. Importantly, desmosomes and keratin filaments influence ER tubule organization and dynamics^10^. However, disrupting desmosomes did not abolish ER localization at cell-cell interfaces, prompting us to investigate additional mechanisms that influence ER organization at intercellular contacts. Furthermore, it is currently unknown whether desmosomes or other adhesive cell-cell junctions influence contacts between the ER and the plasma membrane.

Adherens junctions are adhesion complexes that maintain tissue architecture and regulate epithelial cell morphology and differentiation^15,16^. These junctions require classical cadherins, such as E-cadherin, and are coupled to the actin cytoskeleton through α- and β-catenin^17^. Using a panel of A431 epithelial cells depleted of various adherens junction proteins, we show that E-cadherin association with α-catenin is both necessary and sufficient to recruit ER tubules to cell-cell interfaces. Furthermore, ER recruitment to cell-cell interfaces promotes the formation of ER-PMCS in close spatial and temporal proximity to both adherens junctions and desmosomes. High resolution imaging indicates that adherens junctions, desmosomes, and ER form an adhesion-organelle complex that organizes ER contacts across the plasma membrane. Interestingly, disrupting this complex by ablating adherens junctions altered global cellular lipid content. These findings reveal an unrecognized role for cadherins in organizing ER-PM contacts, providing a means for localized regulation of plasma membrane lipids and architecture.

## Results

### E-cadherin is required for ER localization to cell-cell interfaces

We recently determined that peripheral ER tubules form close, stable associations with desmosomes, and that disrupting desmosomes alters ER tubule dynamics^10^. However, ER tubules still persist at cell-cell interfaces in desmosome-deficient cells, prompting us to investigate whether other intercellular complexes such as adherens junctions regulate ER organization at these sites. To determine if adherens junctions regulate ER localization at cell-cell interfaces, we generated A431 epithelial cells stably expressing a fluorescently tagged pH-stable ER lumenal marker, ER-StayGold, in both wild-type A431 and E/P-cad KO cells in which both E- and P-cadherin genes were ablated^18^. We incubated cells with fluorescently-tagged wheat germ agglutinin (WGA-640R) to visualize the plasma membrane and assessed ER localization by live-cell spinning disk confocal microscopy. In wild-type cells, ER tubules fully extended to cell-cell interfaces (Fig. 1A, top left) but remained more distant from the periphery at the outer edges of multi-cellular colonies (Fig. 1A, arrow). In contrast to wild-type cells, ER tubules failed to extend to cell-cell interfaces in E/P-cad KO cells (Fig. 1A, top right). Re-expression of wildtype E-cadherin in the E/P-cad KO cells (E/P-cadKO+E-cad-WT) restored ER localization to cell-cell interfaces (Fig. 1A, bottom). We quantified ER distribution by measuring the mean fluorescence intensity of ER-StayGold along a radial axis from nucleus to cell edge. In all three cell lines, ER density was highest around the nucleus, where the ER is stacked into sheets. ER density progressively decreased toward the cell periphery, where the ER becomes more tubular. However, in E/P-cad KO cells, this decrease in peripheral ER tubules was more pronounced, and the phenotype was rescued by re-expression of wild-type E-cadherin (E/P-cadKO+E-cad-WT) (Fig. 1B). In the most peripheral region of the cell, E/P-cad KO cells exhibited a two-fold reduction in ER density compared to wild-type cells (Fig. 1C). These findings demonstrate that E-cadherin is required for ER recruitment to cell-cell interfaces.

**Figure 1:**
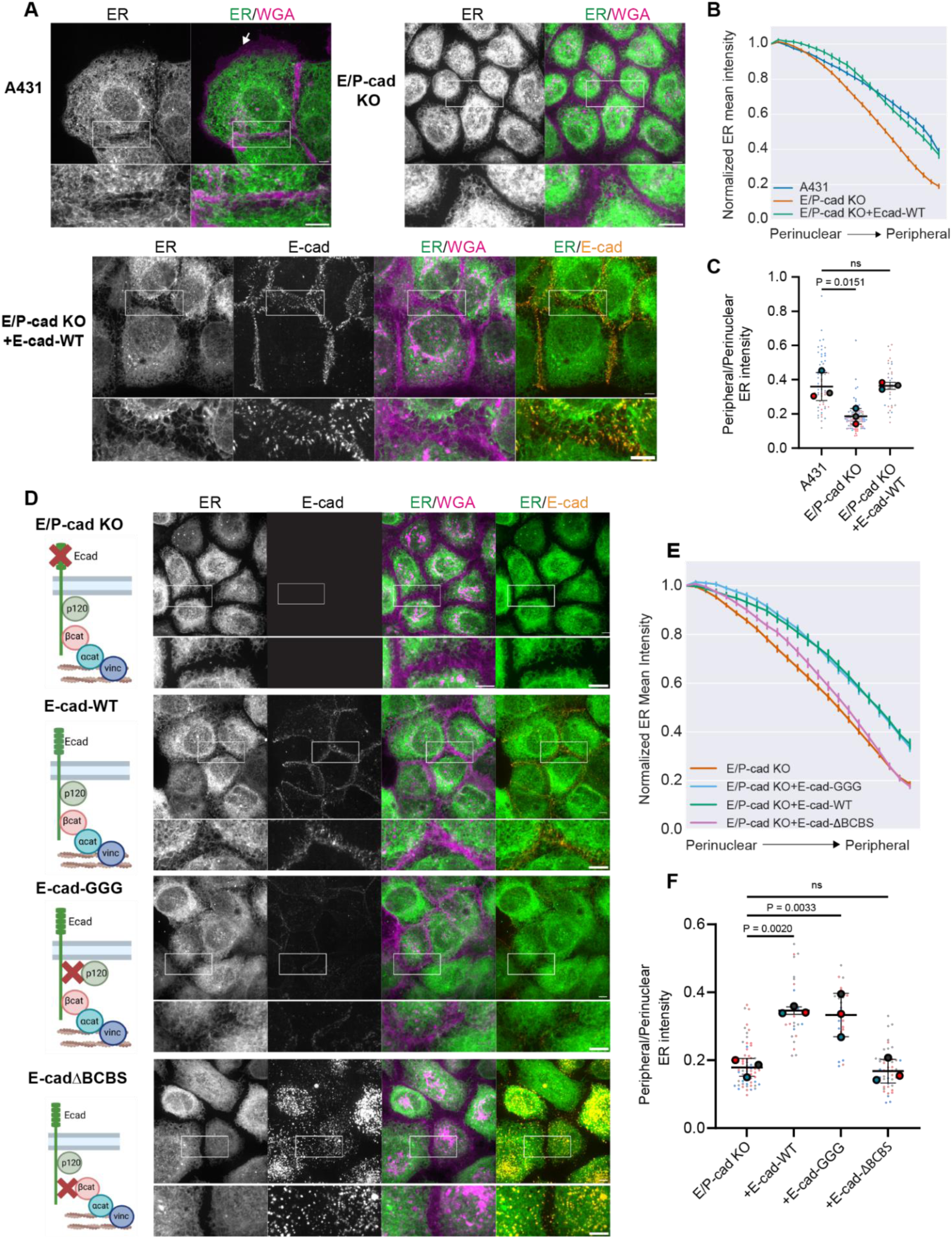
Adherens junctions govern the positioning of ER at cell-cell interfaces. **A**) Live-cell images of wild-type A431 (left), E/P-cad KO (right), and E/P-cad KO+E-cad-WT cells (bottom) expressing ER-StayGold (ER marker, green) and labeled with WGA-640R (plasma membrane marker, magenta). White boxes indicate regions magnified below. Scale bars,10 µm. **B**) Normalized mean fluorescence intensity of ER-StayGold along a radial axis (24 axis points) from nucleus to cell periphery. n=60-100 cells per cell line. Mean ± SD from 3 experiments are shown. **C**) Superplot of the ratio of peripheral to perinuclear ER-StayGold intensity. Small circles = individual cells, large circles = means from 3 experiments. Solid line = overall mean. Error bar = SD. One-way ANOVA with Dunnett’s post hoc test; ns=not significant. **D**) Live-cell images of ER-StayGold, E-cad-RFP, and WGA-640R in E/P-cad KO cells and E/P-cad KO cells expressing different red fluorescent protein (RFP)-tagged E-cadherin mutant proteins. The schematics depict the impact of each E-cadherin mutation on adherens junction. Scale bars, 10 µm **E**) Graph of normalized mean fluorescence intensity of ER-StayGold along a radial axis from nucleus to cell periphery. n=30-80 cells per cell line. Mean ± SD from 3 experiments are shown. **F**) Superplot of the ratio of peripheral to perinuclear ER-StayGold intensity. Solid line = overall mean. Error bar = ± SD. One-way ANOVA with Dunnett’s post hoc test.

We postulated that E-cadherin recruits ER tubules to cell-cell interfaces through association with the microtubule or actin cytoskeleton since both of these networks influence ER organization^19–21^. Adherens junctions associate with microtubules through p120-catenin^22^ and with actin filaments through α-catenin^23,24^. We tested the role of p120-catenin and α-/β-catenin in ER recruitment using E-cadherin mutants deficient in p120-catenin binding (E-cad-GGG) or β-catenin binding (E-cad-ΔBCBS). Interestingly, the expression of E-cad-GGG in E/P-cad KO cells restored ER localization at cell-cell interfaces comparable to E-cad-WT, despite its lower expression level due to increased turnover of the cadherin in the absence of p120-catenin binding^25–28^ (Fig. 1D, 1E). Moreover, the E-cad-GGG mutant associated with both β- and α-catenin and recruited these actin adaptor proteins to cell-cell interfaces (Fig. S1C). In contrast to E-cad-GGG, E-cad-ΔBCBS, which cannot bind β- and α-catenin, failed to localize properly at cell-cell interfaces. Instead, E-cad-ΔBCBS often accumulated as cytoplasmic puncta (Fig. S1), consistent with previous reports of trafficking defects in E-cadherin mutants unable to bind β-catenin^29^. We confirmed by immunofluorescence the lack of p120-catenin and β-catenin association with E-cadherin in E-cad-GGG and E-cad-ΔBCBS cells, respectively (Fig. S1A-B). Quantification of peripheral ER density indicated that E-cadherin accumulation at cell-cell contacts is required for ER recruitment to cell-cell interfaces and that E-cadherin interactions with p120-catenin are dispensable for this function (Fig. 1F).

### Adherens junctions recruit ER to cell-cell interfaces through E-cadherin association with α-catenin

α-catenin stabilizes adherens junctions against mechanical forces by linking E-cadherin to the actin cytoskeleton^30^. To test whether α-catenin is required for ER recruitment to cell-cell interfaces, we imaged ER distribution in A431 cells lacking α-catenin (α-cat KO). Similar to our observations in E/P-cad KO cells, ER tubules were distal from cell-cell interfaces in the α-cat KO cells (Fig. 2A), though E-cadherin and p120-catenin still localized to cell-cell interfaces, albeit at a lower level than in wild-type A431 cells (Fig. S2A). Next, we directly tested the role of E-cadherin association with α-catenin in ER recruitment by expressing an E-cadherin–α-catenin chimeric protein that lacks the binding sites for both p120-catenin and β-catenin (E-cad–αcat)^31^. In addition, we used a variant of this protein, E-cad–αcat-vinc-m, which harbors an α-catenin deletion that eliminates the vinculin binding site. Both α-catenin and vinculin mediate interactions between the cadherin tail and the actin cytoskeleton, with vinculin being particularly important for the ability of adherens junctions to resist mechanical stress^31,32^. We confirmed that vinculin was recruited to adherens junctions with the wild type α-catenin chimera and that this interaction was absent in the E-cad–αcat-vinc-m cells (Fig. S2B). Interestingly, the E-cad–αcat chimera restored ER localization to cell-cell interfaces in E/P-cad KO cells to a level comparable to E-cad-WT (Fig. 2B-D). In contrast, E-cad–αcat-vinc-m failed to fully restore peripheral ER localization. These results suggest that adherens junctions recruit ER to cell-cell interfaces through E-cadherin and α-catenin association, and that this process is partly dependent on vinculin.

**Figure 2:**
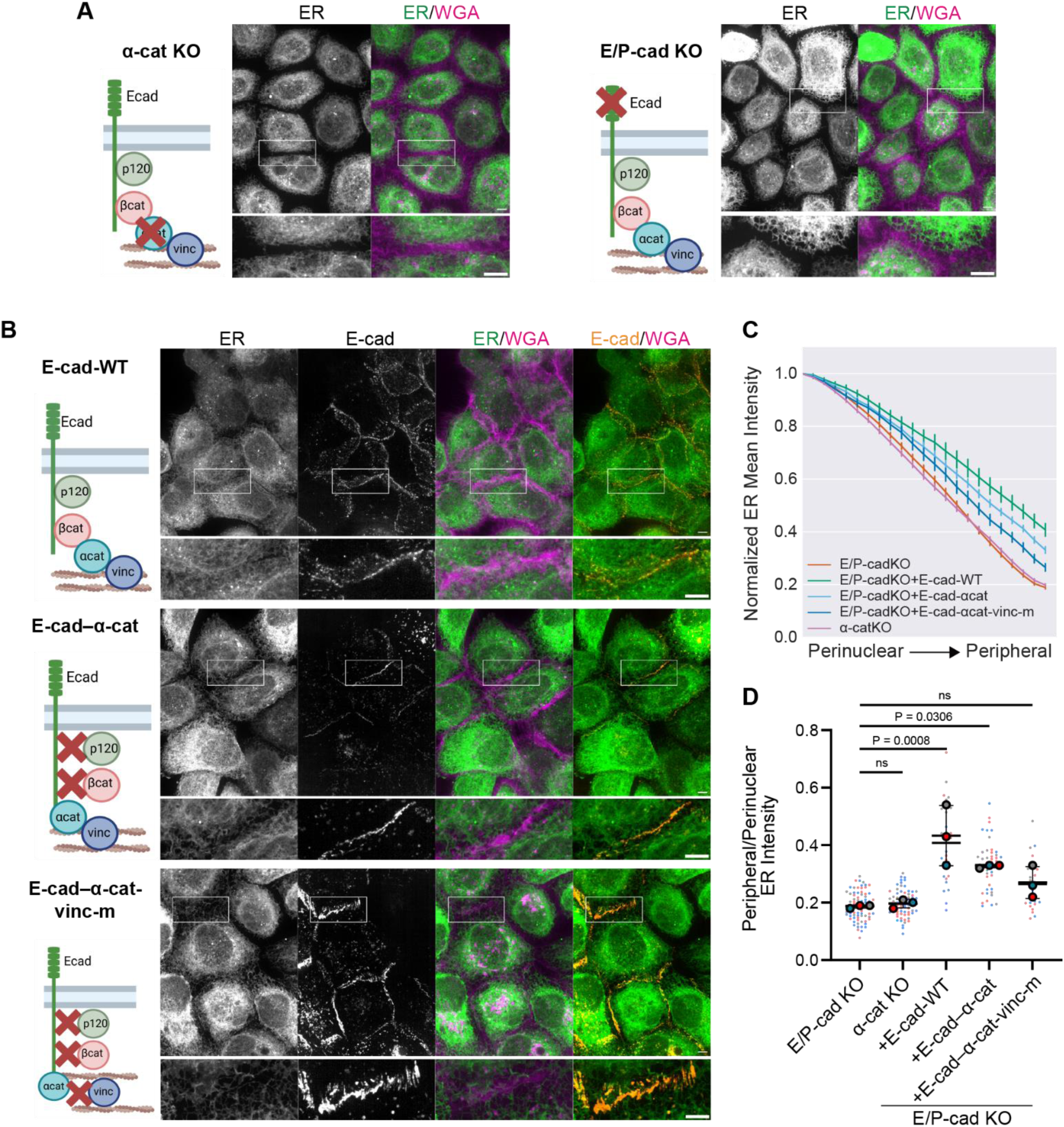
E-cadherin association with α-catenin is required for ER positioning at cell-cell interfaces. **A)** Live-cell images of ER-StayGold in E/P-cad KO cells and α-cat KO cells labeled with WGA-640R. **B**) Live-cell images of ER-StayGold and WGA-640R in E/P-cad KO cells rescued with different RFP-tagged E-cadherin mutant proteins. White boxes indicate regions magnified below. The schematics depict the impact of each E-cadherin mutation on adherens junction. Scale bar, 10 µm. **C**) Graph of normalized mean fluorescence intensity of ER-StayGold along a radial axis from nucleus to cell periphery. n=30-70 cells per cell line. Mean ± SD from 3 experiments are shown. **D**) Superplot of the ratio of peripheral to perinuclear ER-StayGold intensity. Solid line = overall mean. Error bar = ± SD. One-way ANOVA with Dunnett’s post hoc test.

### Adherens junctions and ER appear simultaneously at nascent cell-cell contacts, followed by desmosome formation

To define the temporal hierarchy of ER recruitment to cell-cell interfaces relative to adherens junctions and desmosome formation, we performed a calcium switch assay in A431 cells stably expressing E-cad-HaloTag7 (adherens junction marker), mApple-VAPB (ER marker), and desmoplakin-EGFP (desmosome marker). Cells were initially cultured in low calcium (30 µM) overnight to ablate adherens junctions and desmosomes and subsequently switched to high calcium (2.7 mM) to initiate junction assembly. Immediately after exposure to high calcium, cells were imaged every 60 seconds over a 1.5-hour time course. Initially, ER tubules were distal from the plasma membrane (Fig. 3A, Supplementary Movie 1). Upon cell-cell contact formation, E-cadherin puncta appeared, followed by extension of ER tubules to E-cadherin puncta within 2 minutes (Fig. 3B, F). In most instances, an ER tubule from the adjacent cell extended to the nascent E-cadherin puncta within 5 minutes, forming a mirror image-like arrangement of ER tubules. These events were followed by desmoplakin clustering at regions containing E-cadherin-positive puncta and a paired ER arrangement at cell-cell contacts (Fig. 3D-E). Similar results were obtained using mChilada-α-catenin as an alternative adherens junctions marker (Fig. S3, Supplementary Movie 2). These results support previous studies demonstrating that desmosome assembly occurs after nascent adherens junction formation^33–37^. To resolve the timing of E-cadherin and α-catenin appearance at nascent cell-cell contacts, we imaged A431 cells expressing E-cadherin-mEmerald, mChilada-α-catenin, and HaloTag7-VAPB during calcium-induced junction assembly. In the majority of instances, E-cadherin and α-catenin clustered together at nascent cell-cell contacts (Fig. 3H, L, Supplementary Movie 3). The first ER tubules at nascent cell-cell contacts appeared 2 minutes after E-cadherin and α-catenin (Fig. 3I, L). Often, ER tubules in one cell appeared before ER tubules in the adjacent cell (Fig. 3L). Collectively, these findings indicate that adherens junction assembly is followed closely by ER tubule recruitment. Desmosome assembly is subsequently initiated near adherens junctions at domains where ER tubules have formed paired arrangements at cell-cell contacts.

**Figure 3:**
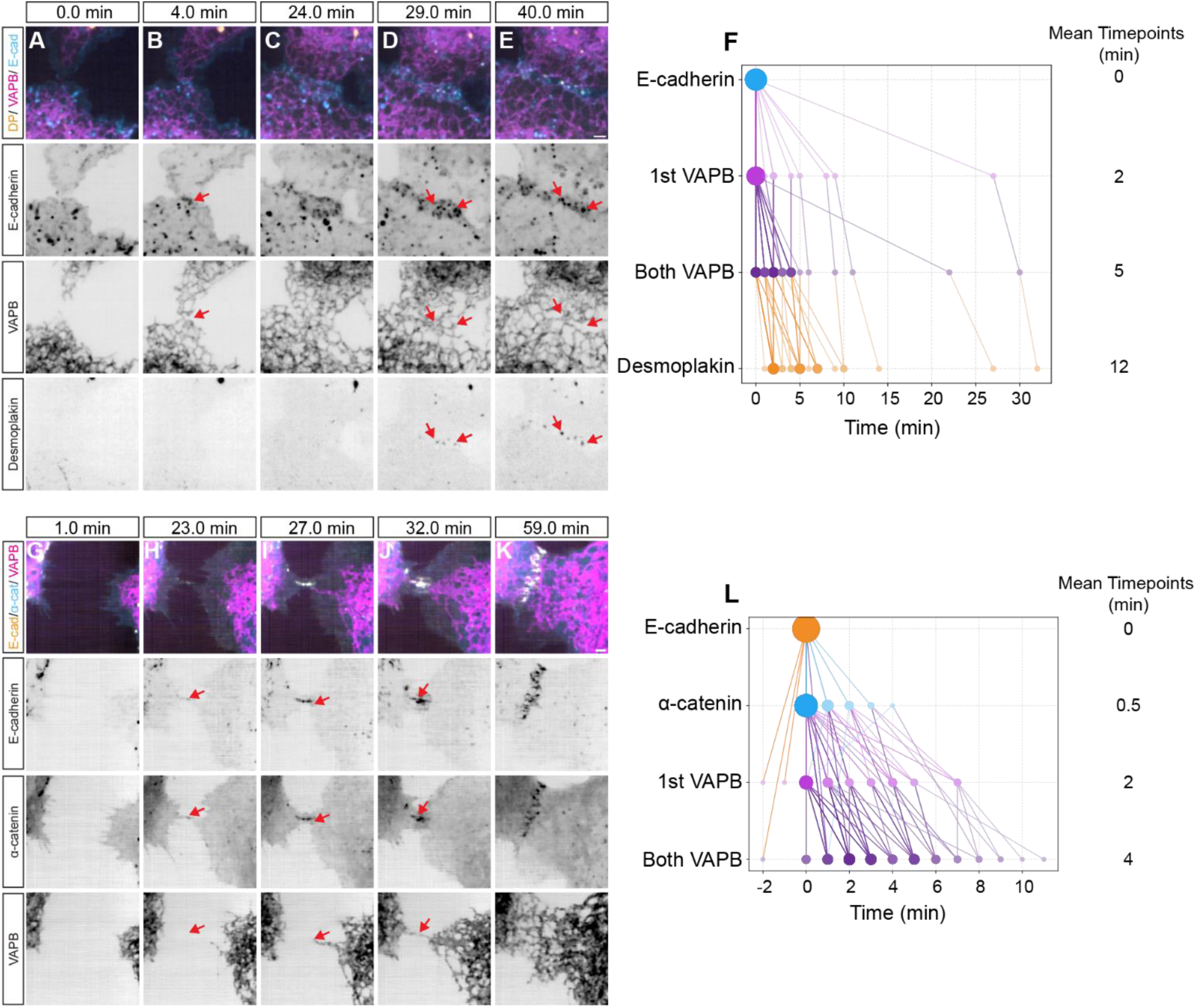
Adherens junctions and ER appear simultaneously at nascent cell-cell contacts, followed by desmosome assembly. A-E) Live-cell, time-lapse images of E-cadherin-HaloTag7 (adherens junction marker), mApple-VAPB (ER marker), and desmoplakin-EGFP (desmosome marker) localization during calcium-induced cell-cell contact formation in A431 cells. **F)** Time course graph indicating the first appearance of E-cadherin, 1st ER (ER in one cell), both ER (ER in both cells), and desmoplakin at nascent cell-cell contacts. Timepoints were subtracted from the timepoint of the first appearance of E-cadherin for each cell-cell contact formation (T=0 for E-cadherin). n=28 cell-cell contact formations. Larger circle size and darker shading corresponds to higher frequency of each protein’s appearance at a given timepoint. **G-K**) Live cell, time-lapse images of E-cadherin, mChilada-α-catenin, and HaloTag7-VAPB during cell-cell contact formation in A431 cells. **L)** Time course graph indicating the first appearance of E-cadherin, α-catenin, 1st ER, and both ER at nascent cell-cell contacts. Timepoints were normalized to the first appearance of E-cadherin for each cell-cell contact formation, as in F. n=70 cell-cell contact formations.

### The ER forms contacts with the plasma membrane adjacent to adherens junctions and desmosomes

The data shown above indicate that adherens junctions recruit ER tubules to cell-cell interfaces. However, it is unknown if ER forms contact sites with the plasma membrane at cell-cell adhesive junctions. We used two complementary approaches to determine if ER tubules form contacts with the plasma membrane at cell-cell junctions. First, we analyzed volume electron microscopy (FIB-SEM) data of A431 cells^10^. Using a deep learning tool (3D U-Net), we segmented the ER, plasma membrane, desmosomes, and ER-PMCS (Fig. 4A-E). We defined ER-PMCS as regions where the ER membrane is within 28 nm of the plasma membrane^38^. ER-PMCS were frequently observed adjacent to desmosomes at cell-cell interfaces (Fig. 4F-L, Supplementary Movie 4), with half of all ER-PMCS being within ∼480 nm of desmosomes (Fig. 4M). In addition, ER-PMCS within 1 µm of desmosomes were larger than those located more distal to desmosomes (Fig. 4N). These data demonstrate that bona fide ER-PMCS localize to plasma membrane domains adjacent to desmosomes at cell-cell interfaces.

**Figure 4:**
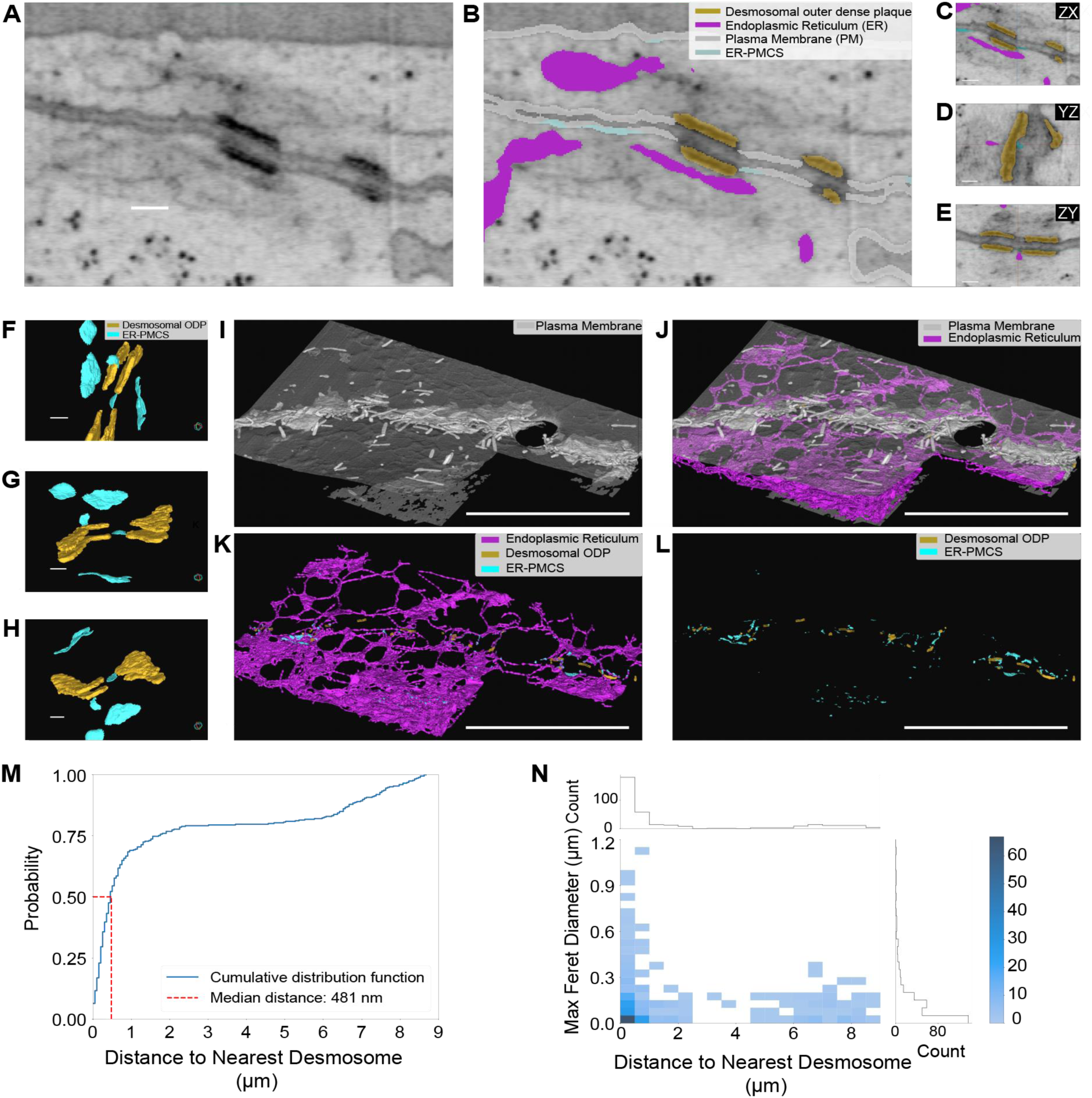
FIB-SEM analysis reveals that ER-PM contact sites are positioned adjacent to desmosomes. **A-B**) 3D FIB-SEM images featuring two desmosomes and nearby ER without segmentations (**A**) and with segmentations of desmosomal outer dense plaque (yellow), ER (magenta), plasma membrane (gray), and ER-PM contact sites (cyan) (**B**). **C-E**) Orthoslices of a region of cell-cell interface where ER-PM contact sites occur adjacent to desmosome. **F-H**) Rotated 3D renderings highlighting ER-PM contact sites (cyan) between two desmosomes (yellow). **I-L**) A cell-cell interface of A431 cells imaged with 3D FIB-SEM and segmented for plasma membrane, ER, desmosomes, and ER-PM contact sites. **I**) The plasma membrane reveals many protrusions at the cell-cell interface. **J**) ER at cell-cell interface are tubular. **K**) ER tubules form ER-PM contact sites at apical and at cell-cell interfaces near desmosomes. **L**) ER-PM contact sites occur near desmosomes. **M**) Cumulative distribution of ER-PM contact sites’ distance to the nearest desmosome. The red dashed line indicates the median distance (481 nm). **N**) 2D histogram showing the relationship between ER-PM contact sites maximum Feret diameter (length) and its distance to the nearest desmosome.The intensity of blue shading in each bin corresponds to the number of contact sites shown in the color bar on the far right. The 1D histogram above the graph depicts the distribution of ER-PM contact sites by distance to nearest desmosome, while the 1D histogram on the right depicts the distribution by maximum Feret diameter. Scalebars, **A-H**) 100 nm **I-L**) 10 μm.

In addition to volume EM analysis, we also performed live-cell imaging of A431 cells to further define the organization of ER and ER-PMCS at cell-cell junctions. We first examined the localization of ER tubules relative to both adherens junctions and desmosomes in A431 cells using super-resolution spinning disk confocal microscopy. Imaging of cells co-expressing mChilada-α-catenin, HaloTag7-VAPB, and DP-EGFP revealed ER tubule tips associating directly with α-catenin-positive puncta at cell-cell interfaces (Fig. 5A, arrows). Moreover, α-catenin and ER tubules often flanked the edges of desmosomes (Fig. 5A, arrow heads). To determine if ER tubules near α-catenin represent ER-PMCS, we used mApple-tagged MAPPER, a well-characterized reporter of ER-PMCS^38^. MAPPER consists of the transmembrane domain of the ER calcium sensor STIM1, which anchors it to the ER membrane, and the polybasic motif of the G-protein Rit, which mediates binding to the plasma membrane when the ER is within 10-25 nm. At cell-cell interfaces, MAPPER organized into both punctate and elongated structures, similar to the morphologies of ER-PMCS observed by volume EM (Fig. 4). Importantly, MAPPER structures localized adjacent to α-catenin and desmoplakin (Fig. 5B). These data indicate that adherens junctions, desmosomes, and ER-PMCS organize into a tripartite assembly at cell-cell interfaces.

**Figure 5:**
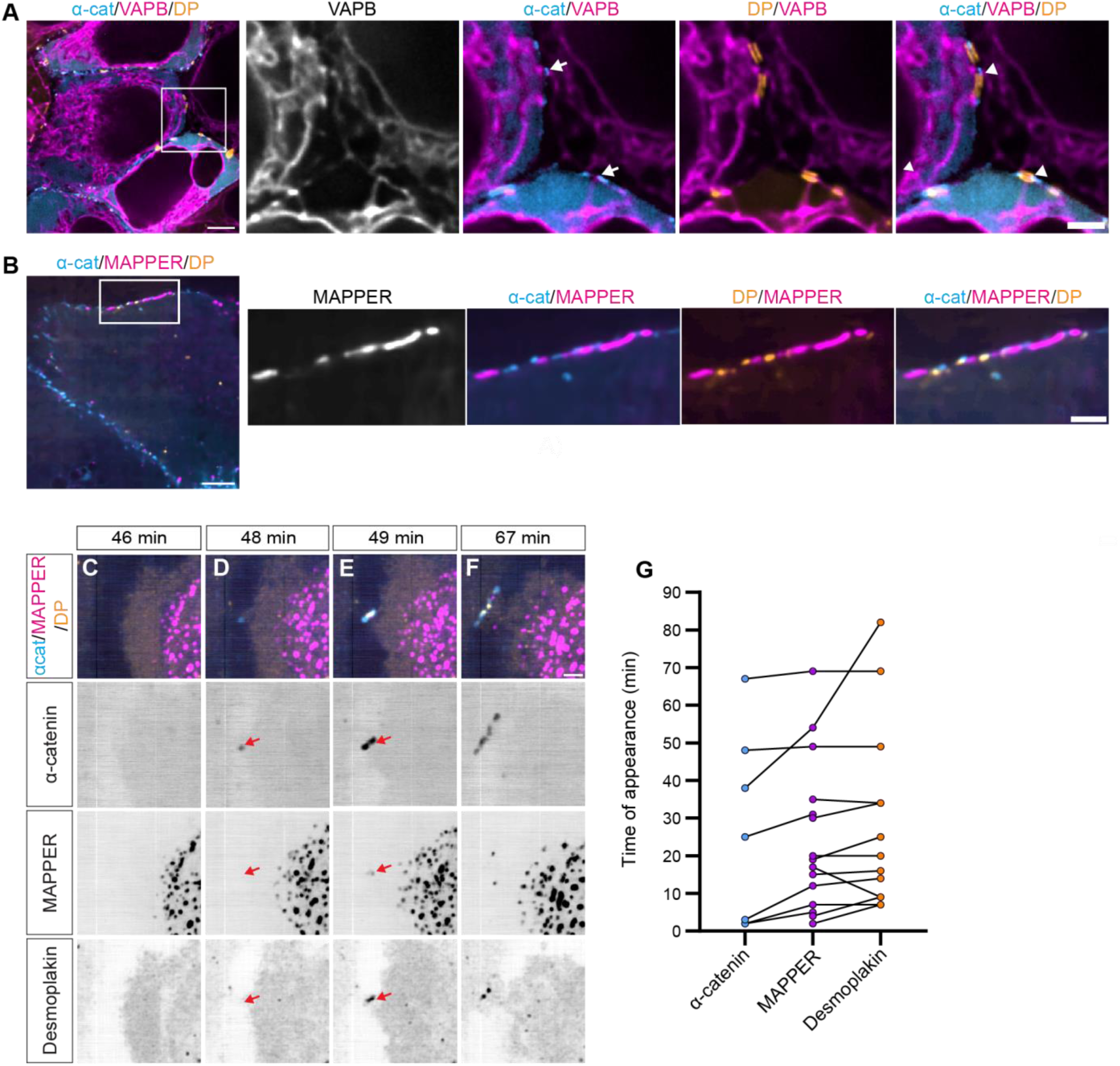
ER-PM contact sites form adjacent to adherens junctions and desmosomes. **A)** Super-resolution, live-cell image of mChilada-α-catenin, HaloTag7-VAPB, and desmoplakin-EGFP in A431 cells. Scale bar, 5 µm. White box indicates region magnified on the right. Scale bar, 2 µm. **B**) Live-cell image of HaloTag7-α-catenin, mApple-MAPPER (ER-PMCS marker), and desmoplakin-EGFP. Scale bar, 5 µm. White box indicates region magnified on the right. Scale bar, 2 µm. **C-F**) Live-cell, time-lapse images of α-catenin, MAPPER, and desmoplakin during calcium-induced cell-cell contact formation. Scale bar, 2 µm. **G**) Time course graph depicting the timepoints at which α-catenin, MAPPER, and desmoplakin first appear at nascent cell-cell contact. Each connected line is one cell-cell contact formation.

To assess the hierarchy of ER-PMCS formation relative to the assembly of adherens junctions and desmosomes, we conducted calcium switch experiments and live-cell imaging of A431 cells co-expressing α-catenin-HaloTag7, mApple-MAPPER, and DP-EGFP. Before cell-cell contact formation, MAPPER puncta were located distal from the leading edge of cells and were predominantly at the cell-substrate interface (Fig. 5C). Adherens junctions formed first, as defined by the appearance of α-catenin puncta at nascent cell-cell contacts (Fig. 5D, arrows), followed by the appearance of MAPPER (Fig. 5E, arrows). On average, MAPPER puncta appeared ∼6 minutes after adherens junctions. Desmoplakin clusters typically appeared simultaneously with or 3 minutes after the formation of MAPPER puncta (Fig. 5E-G). Together, these results indicate that adherens junctions assemble first, followed by ER-PMCS formation and then rapid initiation of desmosome assembly.

### Adherens junctions promote ER-PMCS formation at cell-cell interfaces and regulate cellular lipid levels

The localization and temporal hierarchy of assembly of adherens junctions, desmosomes, and ER-PMCS suggested that adherens junctions might govern the formation of ER-PMCS. To investigate this possibility, we performed live-cell confocal imaging of mApple-MAPPER in wild-type and E/P-cad KO cells. In wild-type cells, numerous MAPPER puncta were present at cell-cell interfaces (3.46/µm border ± 0.36). In contrast, E/P-cad KO cells had fewer MAPPER puncta at cell-cell interfaces (1.65/µm border ± 0.08) (Fig. 6A-B). Quantitative analysis also confirmed a significant reduction in the average MAPPER puncta size at cell-cell interfaces in the E/P-cad KO cells (0.090 µm^2^ ± 0.0095) compared to wild-type cells (0.151 µm^2^ ± 0.022) (Fig. 6B-C). These results indicate that adherens junctions are essential for proper formation of ER-PMCS at cell-cell interfaces.

**Figure 6:**
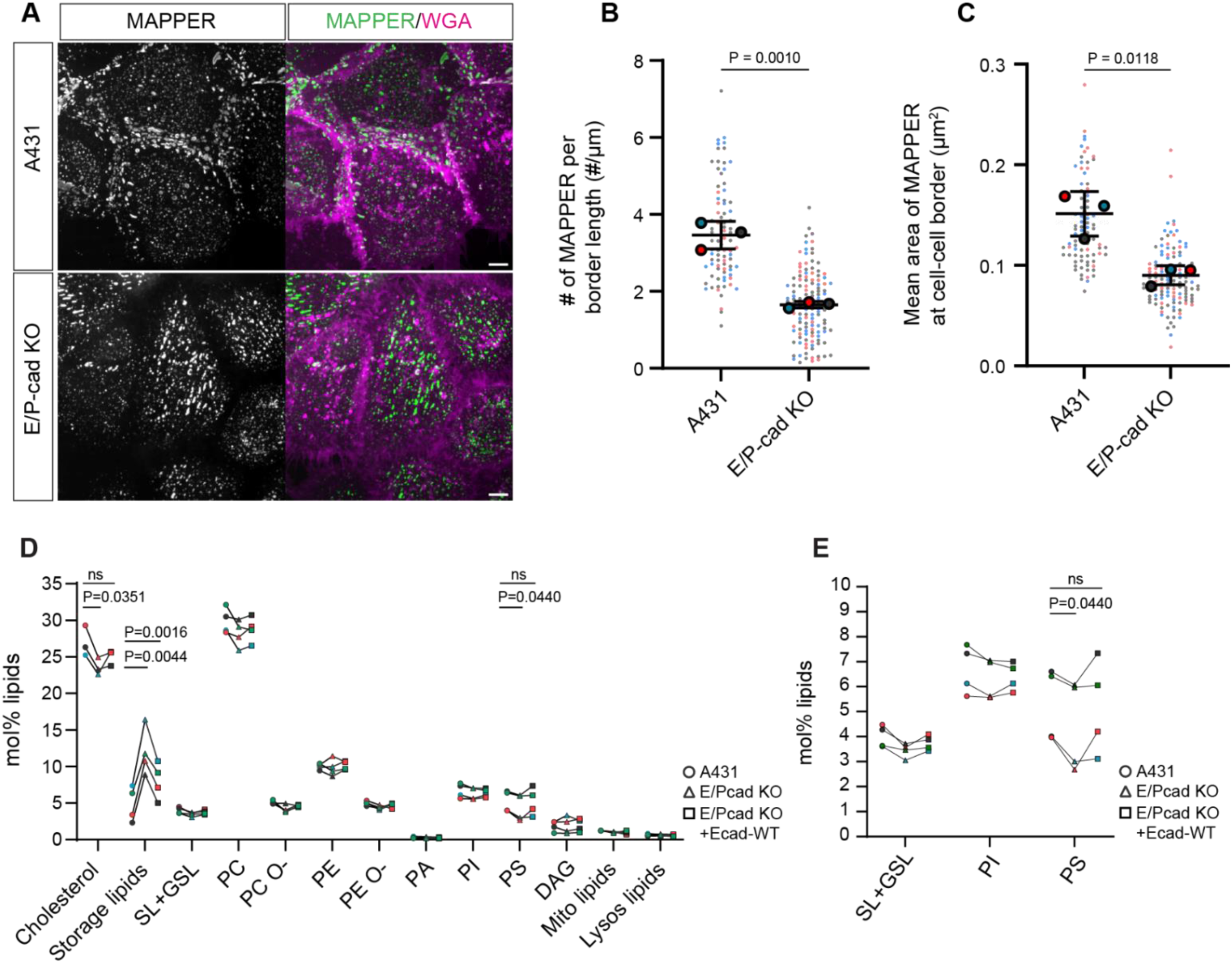
Adherens junctions promote ER-PM contact sites formation at cell-cell interfaces. **A**) Live-cell, maximum intensity projection images of mApple-MAPPER and WGA-640R in wild-type A431 and E/P-cad KO cells. Scale bar, 10 µm. **B**) The number of MAPPER puncta at cell-cell border, normalized to the length of the border measured. **C**) The mean area of MAPPER puncta at cell-cell border. For B and C, dashed line = overall mean ± SD. Two-tailed Student’s t-test. **D)** Molar percentage of each lipid(s) determined by lipidomic analysis of whole cell lysates from wild-type A431, E/P-cad KO, and E/P-cad KO+E-cad-WT cells. Each connected line corresponds to same independent experiment. One-way ANOVA, followed by Dunnett’s test. **E**) Magnified view of selected plasma membrane enriched lipids from D.

Cells rely on both vesicular transport and ER-membrane contact sites to deliver lipids to organelles and the plasma membrane^1,4^. At the plasma membrane, ER-PMCS enable direct exchange of lipids between the ER and plasma membrane. Given that E/P-cad KO cells exhibited fewer and smaller ER-PMCS at cell-cell interfaces compared to wild-type cells (Fig. 6), we next determined whether these differences result in altered cellular lipid composition. We performed mass spectrometry-based lipidomic analysis on whole cell lysates from wild-type, E/P-cad KO, and E/P-cadKO+E-cad-WT cells. Compared to wild-type cells, E/P-cad KO cells showed a marked decrease in several plasma membrane-enriched lipids such as sphingolipids and glycosphingolipids, cholesterol, and phosphatidylserine (PS). Re-expression of E-cad-WT in E/P-cad KO cells partially restored these lipid levels (Fig. 6D-E, Table 1). Conversely, storage lipids, such as triacylglycerol and cholesterol esters, were significantly elevated in the E/P-cad KO cells and rescued by re-expression of wild-type E-cadherin (A431 = 4.9%, E/P-cad KO = 12.0%, E/P-cad KO+E-cad-WT = 8.0%). These findings suggest that adherens junctions regulate cellular lipid composition by regulating ER-PMCS formation and positioning.

**Table 1:**
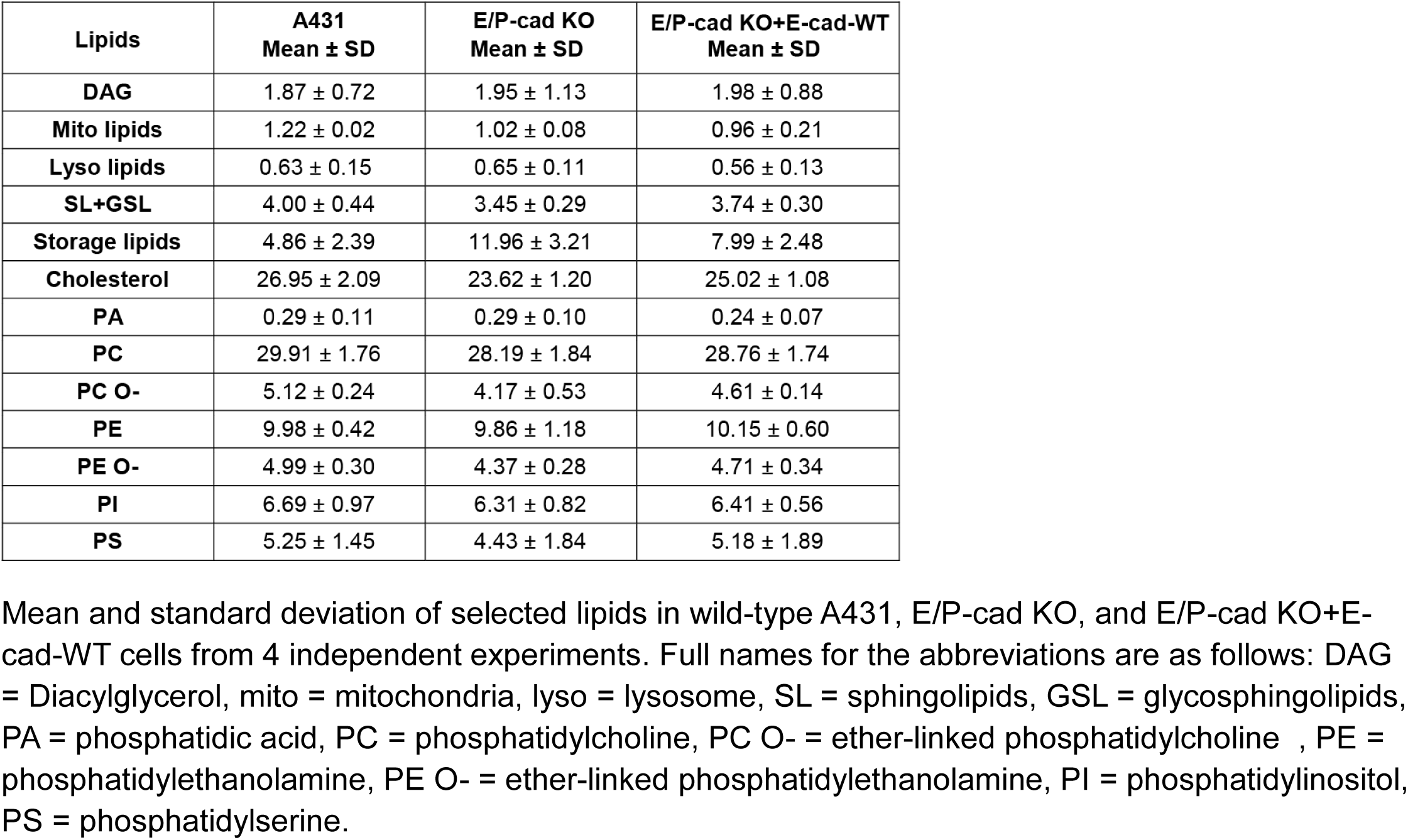
Molar percentage of lipids in whole cell membranes.

| Lipids | A431<br>Mean ± SD | E/P-cad KO<br>Mean ± SD | E/P-cad KO+E-cad-WT<br>Mean ± SD |
| --- | --- | --- | --- |
| DAG | 1.87 ± 0.72 | 1.95 ± 1.13 | 1.98 ± 0.88 |
| Mito lipids | 1.22 ± 0.02 | 1.02 ± 0.08 | 0.96 ± 0.21 |
| Lyso lipids | 0.63 ± 0.15 | 0.65 ± 0.11 | 0.56 ± 0.13 |
| SL+GSL | 4.00 ± 0.44 | 3.45 ± 0.29 | 3.74 ± 0.30 |
| Storage lipids | 4.86 ± 2.39 | 11.96 ± 3.21 | 7.99 ± 2.48 |
| Cholesterol | 26.95 ± 2.09 | 23.62 ± 1.20 | 25.02 ± 1.08 |
| PA | 0.29 ± 0.11 | 0.29 ± 0.10 | 0.24 ± 0.07 |
| PC | 29.91 ± 1.76 | 28.19 ± 1.84 | 28.76 ± 1.74 |
| PC O- | 5.12 ± 0.24 | 4.17 ± 0.53 | 4.61 ± 0.14 |
| PE | 9.98 ± 0.42 | 9.86 ± 1.18 | 10.15 ± 0.60 |
| PE O- | 4.99 ± 0.30 | 4.37 ± 0.28 | 4.71 ± 0.34 |
| PI | 6.69 ± 0.97 | 6.31 ± 0.82 | 6.41 ± 0.56 |
| PS | 5.25 ± 1.45 | 4.43 ± 1.84 | 5.18 ± 1.89 |

Mean and standard deviation of selected lipids in wild-type A431, E/P-cad KO, and E/P-cad KO+E-cad-WT cells from 4 independent experiments. Full names for the abbreviations are as follows: DAG = Diacylglycerol, mito = mitochondria, lyso = lysosome, SL = sphingolipids, GSL = glycosphingolipids, PA = phosphatidic acid, PC = phosphatidylcholine, PC O- = ether-linked phosphatidylcholine, PE = phosphatidylethanolamine, PE O- = ether-linked phosphatidylethanolamine, PI = phosphatidylinositol, PS = phosphatidylserine.

### ER-PMCS predominantly form at sites of cell-cell adhesion

Epithelial cells have polarized plasma membrane domains where the cells adhere to extracellular matrix at the base of the cell and form cell-cell junctions at lateral plasma membrane domains. To assess how adherens junctions influence the spatial distribution of ER-PMCS, we quantified the axial position of individual MAPPER puncta across the entire height of the cell. In z-stack images of mApple-MAPPER-expressing cells labelled with the plasma membrane marker WGA-640R, we classified MAPPER puncta into 3 categories: ‘Basal MAPPER’ were defined as those located within 0.9 µm from the base of the cell; ‘Lateral MAPPER’ were defined as those that colocalized with WGA signal at cell-cell interfaces; and ‘Apical MAPPER’ were defined as those located at the apical domain of the cell. In wild-type cells, basal MAPPER puncta were round and smaller (Fig. 7A), whereas lateral MAPPER puncta were larger and often appeared as linear streaks arranged in a pairwise fashion at both sides of adjacent cells (Fig. 7B, arrows). In contrast to wild-type cells, E/P-cad KO cells displayed more basal MAPPER (Fig. 7D) and fewer lateral MAPPER puncta at cell-cell interfaces (Fig. 7E, arrows). In both wild-type and E/P-cad KO cells, apical MAPPER puncta were substantially less abundant than those at cell-substrate and cell-cell interfaces. We assessed the distribution of MAPPER in each cell line by calculating the percentage of MAPPER at lateral and apical domains as a proportion of total MAPPER puncta. In wild-type cells, 67 % ± 2.8 of MAPPER puncta were lateral/apical whereas in E/P-cad KO cells, only 30% ± 6.2 of MAPPER puncta were lateral/apical (Fig. 7G, H). These data indicate that ER-PMCS preferentially localize to cell-cell interfaces when cell junctions are present. In contrast, loss of desmosomes in A431 cells in which desmoglein-2 is ablated using CRISPR-Cas9 (Dsg null) did not impair MAPPER puncta formation nor their distribution (Fig. S4). Together, these findings demonstrate that adherens junctions regulate the polarized distribution of ER-PMCS across plasma membrane regions.

**Figure 7:**
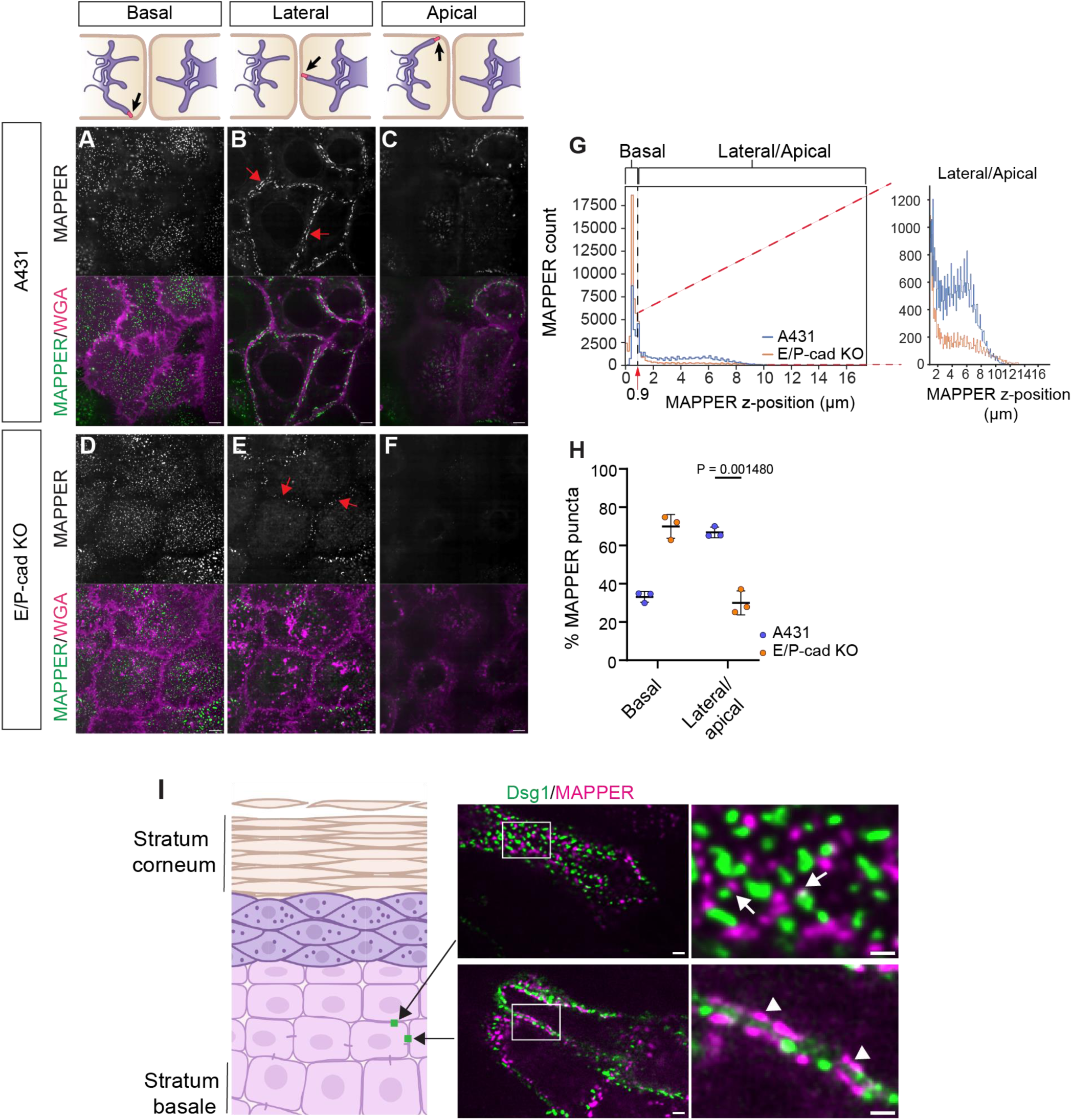
ER-PMCS form at sites of cell-cell adhesion. **A-F**) Single z-plane confocal image of mApple-MAPPER at basal (**A, D**), lateral (**B, E**), and apical (**C, F**) domains in wild-type A431 and E/P-cad KO cells. Schematic at the top of each column indicates plasma membrane domain captured in the images below. Scale bar, 10 µm. **G)** Distribution graph of the number of MAPPER puncta at specific z-positions within cells, with 0 being the base of cell. Dashed line = the transition from basal to latera/apical domains. The graph on the right is a close-up view of the MAPPER distribution at lateral/apical domains. **H**) Percentage of total MAPPER puncta defined as basal or lateral/apical. Mean ± SD from 3 experiments are shown. Solid line = overall mean. Two-tailed Student’s t-test. **I**) The schematic depicts a 3D keratinocyte organotypic culture differentiated into epidermal layers with desmoglein-1 (Dsg1) localization shown in green. Live-cell images of 3D organotypic culture of N/TERT-2G cells expressing Dsg1-GFP and mApple-MAPPER. Scale bar, 2 µm. White boxes indicate regions magnified on the right. Scale bar, 1 µm.

Our data in 2D cell culture systems indicate that ER-PMCS primarily form at regions of the plasma membrane involved in cell-cell and cell-substrate adhesion. We tested this possibility further using an organotypic model of stratifying TERT-immortalized human epidermal keratinocytes (N/TERT-2G)^39^. We transduced these keratinocytes with mApple-MAPPER and Dsg1-GFP to assess ER-PMCS and desmosome localization, respectively. In 3D organotypic cultures, MAPPER localized to cell-cell interfaces adjacent to desmosomes at cell-cell interfaces formed within the same epidermal layer (Fig. 7I, bottom) as well as between basal and suprabasal keratinocytes (Fig.7I, top, arrows). Similar to our observations in 2D cell culture, a mirror image-like arrangement of MAPPER was also observed at lateral cell-cell interfaces (Fig. 7I, bottom, arrow heads). Collectively, these data reveal a structural relationship between adhesive cell-cell junctions and the ER, whereby adherens junctions govern the formation and positioning of contact sites between ER and the plasma membrane.

## Discussion

The spatial organization and dynamics of ER membranes dictate where and when the ER contacts other organelles and the plasma membrane. ER contacts with the plasma membrane regulate lipid and calcium flux to influence a wide range of physiological processes, such as cell migration, cell-matrix adhesion, and plasma membrane repair^5–8^. In this study, we demonstrate that adherens junctions are essential for recruiting the ER to cell-cell interfaces, driven by associations between E-cadherin, α-catenin, and vinculin. ER recruitment to cell-cell interfaces by adherens junctions promotes interactions between the ER and plasma membrane, resulting in the assembly of a tripartite complex comprising ER-PMCS, adherens junctions, and desmosomes. Disruption of adherens junctions destabilizes the organelle–adhesion complex and perturbs cellular lipid homeostasis.

Previous electron microscopy analysis of hepatocytes revealed extensive ER tubule association with cell-cell interfaces, a feature conserved across several epithelial cell models and implicated in regulating hepatocyte differentiation^9^. These findings are consistent with our volume EM studies demonstrating an intimate association of ER tubules with desmosomes^10^. However, the mechanisms regulating ER positioning at cell-cell interfaces remain poorly understood. Our data show that the E-cadherin–α-catenin complex is necessary and sufficient to recruit ER tubules to cell-cell interfaces, whereas cadherin association with p120-catenin is dispensable (Fig. 1D, Fig. 2A, 2B). α-catenin interaction with vinculin further enhances ER recruitment to adhesive cell junctions. For example, an E-cadherin–α-catenin chimeric protein lacking both the p120 binding site and the β-catenin binding domain was sufficient in restoring ER localization to cell-cell interfaces in E- and P-cadherin null cells. However, a variant of this chimeric protein that is deficient in vinculin binding was not fully effective in restoring ER localization to cell-cell interfaces (Fig. 2B). These observations suggest that the link between E-cadherin, the actin cytoskeleton, and actomyosin-generated tension is crucial for ER positioning at cell-cell interfaces. These findings are consistent with a previous study suggesting that ER targeting to adherens junctions depends on local actomyosin tension at adherens junctions^40^.

To further understand the relationship between peripheral ER tubule localization and intercellular junction formation, we determined the spatio-temporal hierarchy of ER recruitment to nascent cell-cell contacts during adherens junction and desmosome assembly. Prior to cell-cell contact formation, ER tubules are positioned distally from lamellipodia (Fig. 3A), similar to ER organization in migrating cells^7^. Upon cell-cell contact formation, ER tubules are rapidly recruited to nascent junctions, often appearing almost simultaneously with adherens junctions, but before desmosome assembly (Fig. 3B-D). As intercellular junctions mature and increase in number, ER tubules progressively accumulate at cell-cell interfaces (Fig. 3E). Lamellipodia contain densely packed and branched actin filaments. Once cells make contact, adherens junctions associate with and reorganize actin filaments through α-catenin^23,41,42^. Thus, actin reorganization during cell junction formation could facilitate ER tubule localization to cell-cell interfaces.

The recruitment of ER tubules to nascent cell-cell interfaces raised the possibility that these tubules interact with plasma membrane domains proximal to adhesive cell junctions. We utilized two complementary approaches to assess ER-PMCS formation at intercellular junctions. Analysis of high-resolution volume EM data (4 x 4 x 4 nm^3^ voxel) revealed that ER-PM contact sites predominantly form adjacent to desmosomes and those contact sites were larger in size than ER-PMCS distal to desmosomes (Fig. 4). These volume EM results were corroborated by optical imaging using a fluorescently tagged reporter of ER-PMCS. ER-PMCS, defined by MAPPER fluorescent puncta, were frequently observed adjacent to both adherens junctions and desmosomes (Fig. 5B). Live-cell imaging data indicate that ER forms contacts with the plasma membrane after adherens junction assembly, followed closely by the appearance of desmoplakin puncta and the initiation of desmosome assembly (Fig. 5C-F). As junction assembly progresses, adherens junction and desmosome components segregate into mature adhesion complexes adjacent to ER-PM contact sites, resulting in a tripartite assembly that mediates both cell-cell adhesion and ER-plasma membrane communication (Fig. 8).

**Figure 8:**
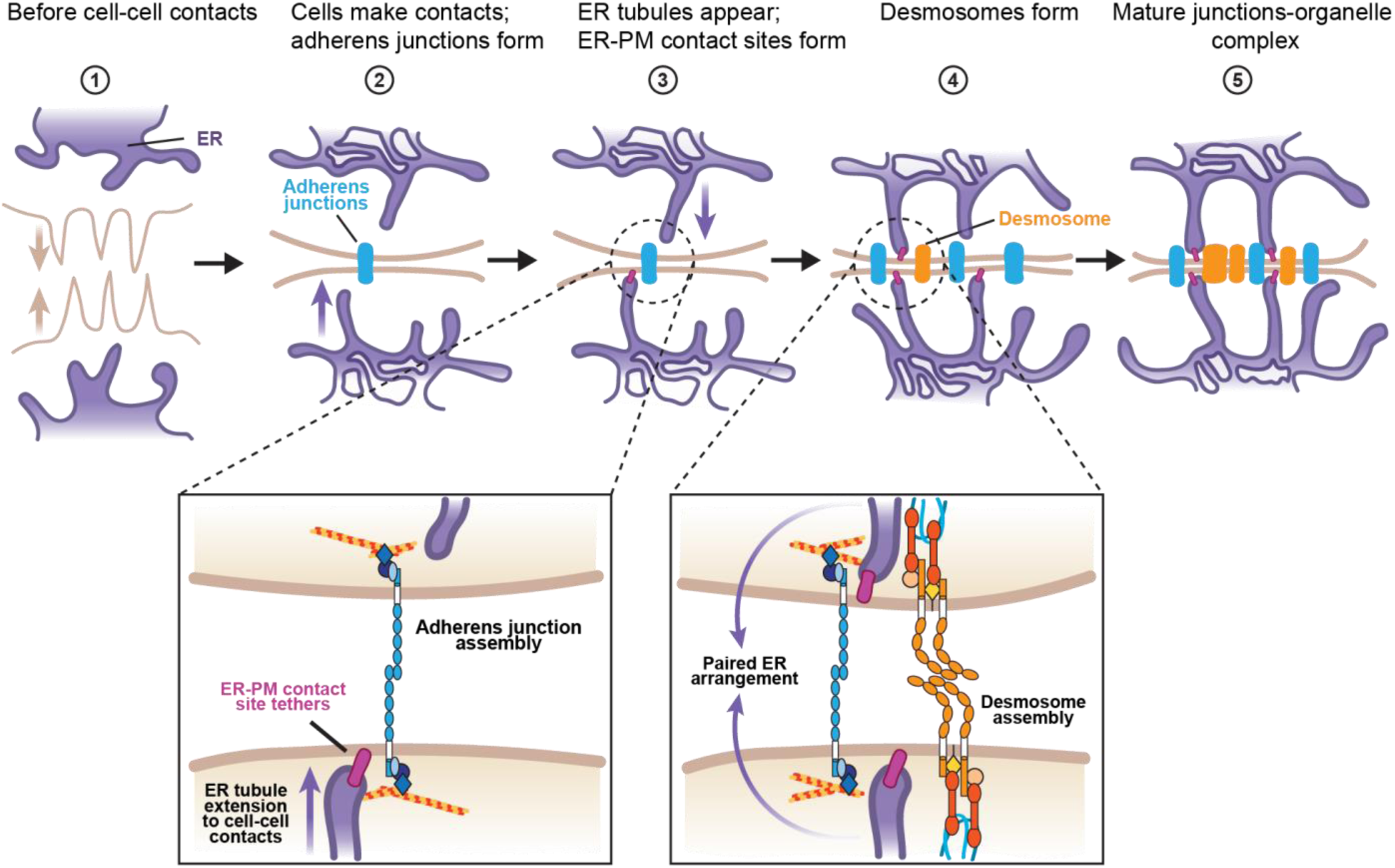
Working model of the stepwise assembly of adherens junction, ER, and desmosome complex at nascent cell-cell contacts. Adherens junctions recruit ER to newly formed cell-cell contacts and promote the formation of ER-PM contact sites. Desmosomes assemble adjacent to adherens junctions at sites of paired ER arrangement. Together, adherens junctions, ER, ER-PMCS, and desmosomes mature into an integrated junction-organelle complex.

This study identifies adhesive cell junctions as a regulator of both the formation and positioning of ER-PMCS. Wild-type A431 cells exhibited more and larger MAPPER puncta at cell-cell interfaces than E/P-cad KO cells (Fig. 6A-C). Moreover, the majority of MAPPER puncta in wild-type cells appeared at cell-cell interfaces, whereas in E/P-cad KO cells, the majority of MAPPER puncta appeared at cell-matrix interfaces (Fig. 7A-H). These results indicate that ER-PMCS preferentially form at plasma membrane regions that contain adhesive cell-cell or cell-matrix contacts. In agreement with this interpretation, ER-PMCS formed at cell-cell interfaces between cells of the basal and suprabasal layers in 3D organotypic epidermal cultures of stratified keratinocytes (Fig. 7I). Collectively, these findings, along with emerging literature, indicate that cell-cell and cell-substrate structures are important regulators of ER distribution and ER contact with the plasma membrane.

ER-PMCS maintain calcium and plasma membrane lipid homeostasis^4,43^. Lipidomic analysis of whole cell membranes from wild-type and E/P-cad KO cells revealed that several lipid species including phosphatidylserine, cholesterol, and sphingolipids were downregulated in E/P-cad KO cells (Fig. 6D-E, Table 1). Reduced cholesterol levels were also observed in an α-catenin knockout epithelial cell line^44^. Interestingly, these lipids are enriched at the plasma membrane and are delivered to the plasma membrane at ER-PMCS^45,46^. In addition, the levels of storage lipids such as cholesterol esters were highly upregulated in E/P-cad KO cells. When cholesterol accumulates in the ER, cells store the excess cholesterol as cholesterol esters inside lipid droplets^47,48^. The decrease in cholesterols and increase in cholesterol esters in E/P-cad KO cells suggests that there is an accumulation of cholesterol in ER, which could be due to defects in cholesterol trafficking from ER to the plasma membrane upon loss of ER-PMCS at cell-cell interfaces. Interestingly, a recent study implicated ORP2 as a lipid transport protein delivering cholesterol from ER to plasma membrane^46^. Together, these results suggest that adherens junctions regulate cellular lipid homeostasis by determining the positioning and size of ER-PMCS.

ER-PMCS localization adjacent to desmosomes in 3D keratinocyte organotypic cultures (Fig. 7) raises the possibility that ER association with cell-cell interfaces could have an important role in tissue homeostasis and architecture. Recent studies indicate that ER-PMCS association with focal adhesions facilitates cell migration by promoting focal adhesion disassembly^5,8^. Cell-cell and cell-matrix adhesive junctions could be influenced by ER regulation of both plasma membrane lipid and/or calcium homeostasis. Interestingly, loss of function in the ER calcium pump sarco-endoplasmic reticulum calcium ATPase 2 (SERCA2) leads to desmosome defects that result in skin fragility^49^. Studies have demonstrated that loss of SERCA2 function impairs desmosome formation by perturbing desmosomal protein trafficking, desmosomal gene expression, and protein kinase C-α signaling^50–53^. Our study raises the possibility that the defect in desmosomes associated with SERCA2 mutations could also be explained by the physical proximity of ER tubules and ER-PMCS with desmosomes and adherens junctions. Furthermore, we recently reported that disrupting desmosomes results in an ER stress response in keratinocytes^10,11^. Collectively, these findings suggest that ER association with the plasma membrane at adhesive junctions integrates cellular mechanics, stress signaling, and lipid metabolism to influence plasma membrane physiology and tissue homeostasis.

## Methods

### Lentivirus generation

Lentivirus transfer plasmids were generated by VectorBuilder except for mApple-VAPB, which was generated in-house as previously described^10^. Vector IDs are listed in Supplementary Table 1 and can be used to retrieve detailed information about the vector on vectorbuilder.com. The sequences of lentivirus transfer plasmids were verified by sequencing. Lentiviruses were made by co-transfecting human embryonic kidney-293FT cells with transfer plasmids, envelope plasmid pMD2.G (encoding VSV-G), and packaging plasmid psPAX1/2 (encoding gag and pol). Supernatants containing lentiviruses were collected 24 h later and for two subsequent days. Lentivirus was concentrated using Lenti-X Concentrator kit (Takara Bio; 631231), followed by centrifugation at 15,000 g at 4°C for 45 minutes. PMD2.G and psPAX2 plasmids (gifts from Didier Trono, plasmid#12259 and #12260) were purchased from Addgene.

### Retrovirus generation and transduction

DSG1-GFP inserted into the pLZRS retroviral vector (a gift from Dr. Kathleen Green, Northwestern University) was transfected into Phoenix 293 cells to produce retroviral supernatants as previously described^54^.

### Cell culture, cell line generation

All cell lines were cultured at 37°C and 5% CO_2_ in a water-jacketed incubator. A431 epithelial cells were cultured in DMEM with L-Glutamine, 4.5g/L glucose, and sodium pyruvate (10013CV; Corning) with 10% fetal bovine serum (TET tested, S10350; R&D Systems) and 1x antibiotic-antimycotic solution (30-004-CI; Corning).

Murine fibroblast (J2-3T3, male) was a gift from Dr. Kathleen Green Lab, Northwestern Univ., Chicago, IL. They were cultured in DMEM with 4.5 g/L glucose (Corning #10–017-CM), 2 mM GlutaMAX (GIBCO #35050061), 100 U/mL penicillin plus 100 mg/mL streptomycin (Sigma #P0781), and 10% fetal bovine serum (FBS; HyClone #SH3007103). J2-3T3 cells were utilized only as supporting cells for organotypic epidermal cultures, and their identity was not independently authenticated. TERT-immortalized male human epidermal keratinocytes N/TERT-2G^39^ were propagated in keratinocyte serum-free medium (KSFM, GIBCO #17005042, supplemented with 30 μg/ml bovine pituitary extract, 0.2 ng/ml epidermal growth factor, 0.31 μM calcium chloride) containing 100 U/mL penicillin plus 100 mg/mL streptomycin (Sigma #P0781). N/TERT-2G were authenticated by their ability to differentiate into a stratified epidermis; no other identity authentication was performed.

A431 E-/P-cadherin CRISPR knockout and A431 α-catenin CRISPR knockout were gifts from Sergey Troyanovsky^18^. A431 cells stably expressing fluorescently-tagged proteins were generated by lentivirus transduction. Cells were incubated with cell culture media containing lentivirus and 10 ug/ml polybrene (TR-1003-G; EMD Millipore) for 1 day and then cultured for another 3 days before selecting for cells stably expressing lentivirus construct. Cells were selected with either blasticidin (5 ug/ml) (R21001; Thermofisher) or puromycin (0.5 ug/ml) (ABT-440; Boston BioProducts) or both depending on the lentivirus construct used. Cells expressing fluorescently-tagged proteins were enriched using fluorescence-activated cell sorting.

For DSG1-GFP retrovirus transduction of N/TERT-2G cells, the cells were cultured in retrovirus-containing medium supplemented with 4 µg/ml polybrene for 60 minutes at 37°C. Cells were then rinsed in PBS and re-fed in their native medium and passaged upon reaching 70%-80% confluency. Expression of fluorophore-tagged viral constructs was visible after 24 hrs.

### Organotypic epidermal culture

Human organotypic epidermal cultures were generated as previously described^55^. Organotypic cultures were generated from TERT-immortalized human epidermal keratinocytes N/TERT-2G^39^ virally transduced with Dsg1-GFP and mApple-MAPPER and differentiated at an air-medium interface using CnT-Prime Epithelial 3D Airlift Medium (CELLnTEC #cnT-PR-3D) atop J2-3T3 fibroblast-populated type 1 collagen rafts.

To make fibroblast-collagen rafts, J2-3T3 fibroblasts were trypsinized, re-suspended in DMEM+10% FBS, counted, centrifuged for 5 min at 200 g, and the supernatant was aspirated. The cell pellet was re-suspended in 1/10 the final volume required (2 mL per organotypic culture) using 10X reconstitution buffer (1.1 g of NaHCO3 plus 2.39 g of HEPES in 50 mL of 0.05 N NaOH) and 1/10 the final volume of 10X DMEM (Sigma #D2554) was then added. High-concentration rat tail type 1 collagen (Corning #354249) was added to achieve a final concentration of 4 mg/ml with sterile water added to adjust to the final volume. The collagen/fibroblast slurry was mixed by several inversions, then pipetted into the top chamber of a transwell insert (Falcon #353091) housed within a deep 6-well culture plate (Falcon #355467). The slurry was allowed to solidify in a tissue culture incubator at 37°C for 60 min. The polymerized fibroblast rafts were submerged in DMEM+10% FBS and incubated at 37°C for at least 24 hr.

Next, N/TERT-2G cells were trypsinized and re-suspended in DMEM+10% FBS. Cells were counted and the required volume was centrifuged at 200 g for 5 min. The supernatant was aspirated and the cell pellet was re-suspended in DMEM+10% FBS to a final volume of 2 mL per organotypic culture. The medium was aspirated from both the upper and lower chambers of the transwell without disturbing the collagen rafts. N/TERT-2G cells were seeded atop the raft (2 million cells) and DMEM+10% FBS was added to the bottom chamber to submerge the insert and raft; cultures then were incubated at 37°C for 24-48 hr at which point the medium was removed from both chambers. Finally, 10 mL of 3D Airlift Medium was added only to the bottom chamber of the transwell, reaching the underside of the raft and exposing the overlying N/TERT-2G cell monolayer to the air. Cultures were fed 3D Airlift Medium every other day for 7 days prior to performing live spinning disk confocal imaging.

### Spinning disk microscope

Fluorescence imaging was performed using a Nikon Ti2-E equipped with a Yokogawa CSU-X1 (or W1) spinning disk unit, LUNF XL laser unit, Nikon Perfect Focus System, Z piezo stage, motorized XY stage, two sCMOS cameras (ORCA-Fusion BT, Hamamatsu Corp.), and two fast filter wheels with elements controlled through hardware-triggering through Nikon’s National Instruments Breakout Box. The acquisition software was NIS-Elements (v5.30.02, v5.30.04, v5.30.06, and 5.42.03). The polychroic mirror within the Yokogawa CSU-X1 unit is a Semrock Di01-T405/488/568/647. Single-emission filters (Chroma ET525/36m, Chroma ET605/52m, and Chroma ET705/62m) were used with the 488 nm, 561 nm, and 640 nm lasers.

### WGA labeling

To visualize plasma membranes, cells were incubated with 2-4 ug/ml WGA-640R (Biotium; 29026) in 1X HBSS with calcium and magnesium (HBSS+) or 1X HBSS without calcium and magnesium (HBSS-) (for calcium switch experiments) at 37°C for 10 minutes. Cells were then washed with 1X HBSS+ or 1X HBSS-. For live-cell imaging, cells were incubated with pre-warmed live-cell imaging media before imaging. For fixed-cell imaging, cells were fixed immediately.

### Live-cell imaging

Cells were seeded on an 8-well chambered #1.5H cover glass (#C8-1.5H-N; Cellvis). Nikon 100x/1.49 NA Apo TIRF oil immersion objective with its correction collar optimized for imaging at 37°C through a #1.5H glass coverslip was used. Images were taken in 12-bit with high gain (‘12-bit Sensitive’) and with ‘Standard’ readout mode. Coarse alignment for simultaneous dual-channel imaging was achieved using spinning disk pinhole alignment until pixel-perfect overlap was achieved in the center of field of view for the two cameras. Physiological temperature and CO_2_ levels (5%) were achieved with Tokai Hit stage top incubation system (model STXF-WELSX-SET).

For confocal imaging of live organotypic epidermal cultures, N/TERT-2G transduced with fluorophore-tagged constructs were seeded onto fibroblast-collagen rafts. After 7 days, stratified cultures were prepared for imaging by removing the entire culture from the transwell using a scalpel and transferring into a 35 mm glass-bottom dish (MatTek #P35G-1.5–20-C). Images of live organotypic epidermis were acquired on a Hamamatsu ORCA-FusionBT sCMOS camera using a Yokogawa CSU-W1 spinning disk confocal system on a Nikon Ti2 microscope. Samples were illuminated using 488, 561, 640 nm laser excitation lines, and fluorescence was detected using a 60× 1.2 NA water objective (Nikon) with standard emission filters. Multi-color Z stack images were acquired using a piezo motor with a step size of 200 nm.

### Indirect immunofluorescence

Cells were grown to at least 50% confluency on #1.5H glass coverslips in tissue culture plates. Cells were fixed in 4% paraformaldehyde (PFA) in 1X PBS with calcium and magnesium (PBS+) for 10 minutes at room temperature and washed 3X with 1x PBS+. For staining of adherens junction and desmosomal proteins, cells were permeabilized and blocked in 1X PBS+ containing 0.1% Triton and 1% BSA for 15 minutes. Cells were incubated in primary antibodies in blocking buffer (1X PBS+ containing 0.1% Triton and 1% BSA) for 1 hour at room temperature, followed by 3 washes in 0.1% Triton in 1X PBS+. Cells were then incubated in secondary antibodies in blocking buffer for 45 minutes at room temperature, followed by 3 washes in 0.1% Triton in 1X PBS+. Coverslips were mounted onto glass microscope slide using ProLong Glass mounting medium (Thermo Fisher Scientific; P36980) and cured overnight.

### Super-resolution imaging system

Super-resolution live-cell images of α-catenin, VAPB, and desmoplakin were acquired on a Nikon Ti2-E microscope equipped with a Yokogawa CSU-W1 spinning disk unit in super-resolution (SoRa) mode. Cells were maintained at 37°C and 5% CO_2_ with the Tokai Hit stage top incubation system. Images were taken with a 60X/1.49 NA Apo TIRF oil objective with 2.8x increased magnification and were deconvolved in NIS elements.

### Image processing

Live-cell snapshots of ER-StayGold were denoised using Denoise.ai software on NIS-Elements. Calcium switch images were denoised using Noise2Void v0.2.1 (N2V), a self-supervised deep learning algorithm. Denoising was performed on single channel and single timepoint images. After re-combining into a hyperstack, all datasets were corrected for lateral chromatic aberration using NanoJ’s channel registration. N2V was used with a TensorFlow-DirectML on a Windows10 workstation with an NVIDIA RTX 3090 GPU.

ImageJ was used to process images for making montages. Images were adjusted for brightness and contrast using lookup tables (LUTs), split by channels and timepoints, and displayed as montages.

### Quantification of ER distribution

Quantification of ER distribution was done on maximum intensity projection images. Images were flat field corrected to remove non-uniform illumination. Regions of interest (ROIs) were manually drawn on the plasma membrane and nuclear borders based on WGA-640R and ER-StayGold labeling, respectively. Each cell was divided into 24 concentric rings from the nucleus to the plasma membrane. The mean fluorescence intensity of ER-StayGold in each ring was divided by the mean fluorescence intensity of ER-StayGold in the perinuclear ring to normalize for differences in ER-StayGold intensities between different cells and biological replicates. To batch process images with saved ROIs, Andrew Moore’s ImageJ macro for radial interpolation was modified (https://github.com/andmoo91/Radial-Intensity-Analysis). Data wrangling was performed using panda (2.2.1) and Python. Data plotting was performed using seaborn (0.13.2) and matplotlib (3.8.3).

### Quantitative analysis of MAPPER puncta

Quantification of number and area of individual MAPPER puncta at cell-cell borders was done on maximum intensity projection images that were denoised using Denoise.ai software on NIS-Elements. MAPPER images were segmented using the Pixel Classification workflow in ilastik. Briefly, features were drawn to distinguish background from foreground (MAPPER). The resulting binary images were overlaid with the corresponding WGA-640R fluorescence images. ROIs were manually drawn on WGA-labeled cell-cell borders. An in-house ImageJ macro was used to quantify the number and area of MAPPER puncta within the ROIs, and the area and maximum Feret diameter (as a proxy for length) of individual ROIs. The number of MAPPER puncta at cell-cell borders was calculated by dividing the total number of MAPPER puncta in each ROI by the length of the ROI. The mean area of MAPPER puncta was calculated by dividing the total area of all MAPPER puncta in each ROI by the number of MAPPER puncta.

To measure the axial/Z-axis position of MAPPER puncta, Z-stack images of MAPPER were segmented using the 3D Pixel Classification workflow in ilastik and processed by an in-house ImageJ macro. 26-connectivity was used to create separate 3D objects, and centroid positions were measured using MorphoLibJ (v 1.6.4).

### ER-PM contact analysis in FIB-SEM data

ER-PMCS analysis on FIB-SEM data with 4 x 4 x 4 nm^3^ voxels focused on cell-cell interfaces between A431 epithelial cells due to time consuming requirements of 3D FIB-SEM. 3D segmentations of ER, plasma membrane, and desmosomes were performed using a 3D U-Net, configured by nnU-Net v2.2 combined with human proofreading^56^. Defining an ER-PMCS as a region of the PM where the ER is within 28 nm, we dilated the ER segmentation and applied a 26-connected component analysis to the union of PM and dilated ER to generate distinct objects using Dragonfly software, Version 2024.1 for Windows.

### Lipidomics of whole cell membranes

Cells were cultured to 95% confluency and washed 3x with cold 1X PBS without Ca^2+^ and Mg^2+^ (PBS-). All the subsequent steps for homogenizing cells were done on ice or at 4°C. Cells were scraped in 1ml 1X PBS-using cell scraper and collected into Eppendorf tubes. Cells were pelleted by centrifugation at 200 rcf for 5 minutes, and pellets were washed with 1X PBS-. Centrifugation and wash were repeated 1x. The cell pellet was resuspended in 300 µl 1X PBS- and the cell number was counted using hemacytometer in order to obtain similar number of cells for lipid comparisons between different cell lines. Cell concentration of 3000 cells/µl were prepared in 400 µl of 1X PBS-. Cells were then homogenized with 25 strokes of 25G 5/8 needle, fast-frozen in liquid nitrogen, and their detailed lipid compositions were analyzed by shotgun electron spray ionization with tandem MS-MS (ESI-MS/MS) by Lipotype, GmbH (Dresden, Germany) and detailed below.

### Lipid extraction for mass spectrometry lipidomics

Mass spectrometry-based lipid analysis was performed by Lipotype GmbH (Dresden, Germany) as described. Lipids were extracted using a two-step chloroform/methanol procedure. Gangliosides were extracted from the water phase of the preceding chloroform/methanol extraction with a solid phase extraction protocol. Samples were spiked with internal lipid standard mixture containing: cardiolipin 14:0/14:0/14:0/14:0 (CL), ceramide 18:1;2/17:0 (Cer), diacylglycerol 17:0/17:0 (DAG), hexosylceramide 18:1;2/12:0 (HexCer), dihexosylceramide 18:1;2/12:0 (DiHexCer), Globoside 3 18:1;2/17:0 (Gb3), GM3-D3 18:1;2/18:0 (GM3), GM1-D3 18:1;2/18:0 (GM1), lyso-phosphatidate 17:0 (LPA), lyso-phosphatidylcholine 12:0 (LPC), lyso-phosphatidylethanolamine 17:1 (LPE), lyso-phosphatidylglycerol 17:1 (LPG), lyso-phosphatidylinositol 17:1 (LPI), lyso-phosphatidylserine 17:1 (LPS), phosphatidate 17:0/17:0 (PA), phosphatidylcholine 15:0/18:1 D7 (PC), phosphatidylethanolamine 17:0/17:0 (PE), phosphatidylglycerol 17:0/17:0 (PG), phosphatidylinositol 16:0/16:0 (PI), phosphatidylserine 17:0/17:0 (PS), cholesterol ester 16:0 D7 (CE), sphingomyelin 18:1;2/12:0;0 (SM), sulfatide d18:1;2/12:0;0 (Sulf), triacylglycerol 17:0/17:0/17:0 (TAG) and cholesterol D6 (Chol). Gb4 was estimated semi-quantitatively, using the Gb3 standard. After extraction, the organic phase was transferred to an infusion plate and dried in a speed vacuum concentrator. 1st step dry extract was re-suspended in 7.5 mM ammonium acetate in chloroform/methanol/propanol (1:2:4; V:V:V) and 2nd step dry extract in 33% ethanol solution of methylamine in chloroform/methanol (0.003:5:1; V:V:V). All liquid handling steps were performed using Hamilton Robotics STARlet robotic platform with the Anti Droplet Control feature for organic solvents pipetting.

### MS data acquisition

Samples were analyzed by direct infusion on a QExactive mass spectrometer (Thermo Scientific) equipped with a TriVersa NanoMate ion source (Advion Biosciences). Samples were analyzed in both positive and negative ion modes with a resolution of R_m/z=200_=280000 for MS and R_m/z=200_=17500 for MSMS experiments, in a single acquisition. MSMS was triggered by an inclusion list encompassing corresponding MS mass ranges scanned in 1 Da increments. Both MS and MSMS data were combined to monitor CE, Chol, DAG and TAG ions as ammonium adducts; LPC, LPC O-, PC, PC O-, as formiate adducts; and CL, LPS, PA, PE, PE O-, PG, PI and PS as deprotonated anions. MS only was used to monitor LPA, LPE, LPE O-, LPG and LPI as deprotonated anions; Cer, HexCer, and SM as formiate adducts and cholesterol as ammonium adduct of an acetylated derivative.

### Data analysis and post-processing

Data were analyzed with in-house developed lipid identification software based on LipidXplorer. Data post-processing and normalization were performed using an in-house developed data management system. Only lipid identifications with a signal-to-noise ratio >5 and a signal intensity 5-fold higher than in corresponding blank samples were considered for further data analysis.

The following lipid names and abbreviations are used: ceramide (Cer), cholesterol, sphingolipids (SL), glycosphingolipids (GSL), diacylglycerol (DAG), phosphatidic acid (PA), phosphatidylcholine (PC), phosphatidylethanolamine (PE), phosphatidylinositol (PI), phosphatidylserine (PS), and their respective lysospecies (lysoPA, lysoPC, lysoPE, lysoPI, and lysoPS) and ether derivatives (PC O-, PE O-, LPC O-, and LPE O-).

## Quantification and statistical analysis

For quantitative analysis of ER distribution and MAPPER puncta, 7-14 images were taken per cell line, per replicate for 3 independent replicates. Images were flat field corrected to remove effects of non-uniform illumination before quantification. Quantifications were performed using ImageJ macros. Statistics were calculated using GraphPad Prism 10.5.0. Statistical significance was measured using one-way ANOVA, followed by Dunnett’s post-hoc test or two-tailed T-test. P-values are reported in figure legends. Superplots were created to depict variations within the same population and between independent experiments.

For calcium switch, live-cell imaging experiments, 3-6 movies were taken per replicate for ≥ 3 independent replicates. Movies were denoised using the Noise2Void algorithm. The time at which adherens junction, ER, and desmosomal proteins first appear at newly forming cell-cell interfaces was determined by the appearance of fluorescence signal that remained at cell-cell interfaces as junctions matured. For some of the calcium switch imaging experiments, only two of the three channels were acquired due to difficulty in finding cells that co-express all three fluorescently-tagged proteins. Lipidomics of whole cell membranes were performed using a similar number of cells for all 3 independent replicates. FIB-SEM data were limited to a one cell-cell interface due to difficulties in sample preparation and image processing. Super-resolution spinning disk confocal data were limited to one replicate that was acquired during a Nikon CSU-W1 SoRa demonstration.

## Data availability

All code, macros, and plasmid maps are available at: https://github.com/KowalczykLab/ER-AJ-manuscript-SL-NKB.

## Acknowledgements

We thank Sergey M. Troyanovsky for providing wild-type, E/P-cad KO, and α-catenin KO A431 cells. We thank Nathan C. Shaner and Gerard G. Lambert (University of California San Diego School of Medicine) for providing the mChilada sequence. We thank Fred Murphy and Lauren Field for images acquired using a Nikon CSU-W1 SoRa microscope. We thank J. Bednarczyk and staff from Penn State College of Medicine’s Flow Cytometry Core for assistance with cell sorting. This work was supported by: National Institutes of Health grant R01AR048266 (A.P.K.); Children’s Miracle Network Faculty Research grant 1110000016 (N.K.B.). Cryo-SIM and FIB-SEM imaging were done in collaboration with the Advanced Imaging Center at Janelia Research Campus, a facility jointly supported by the Gordon and Betty Moore Foundation and the Howard Hughes Medical Institute.

