## Supplemental Figures for "Cadherin Regulation of Endoplasmic Reticulum-Plasma Membrane Contact Sites"

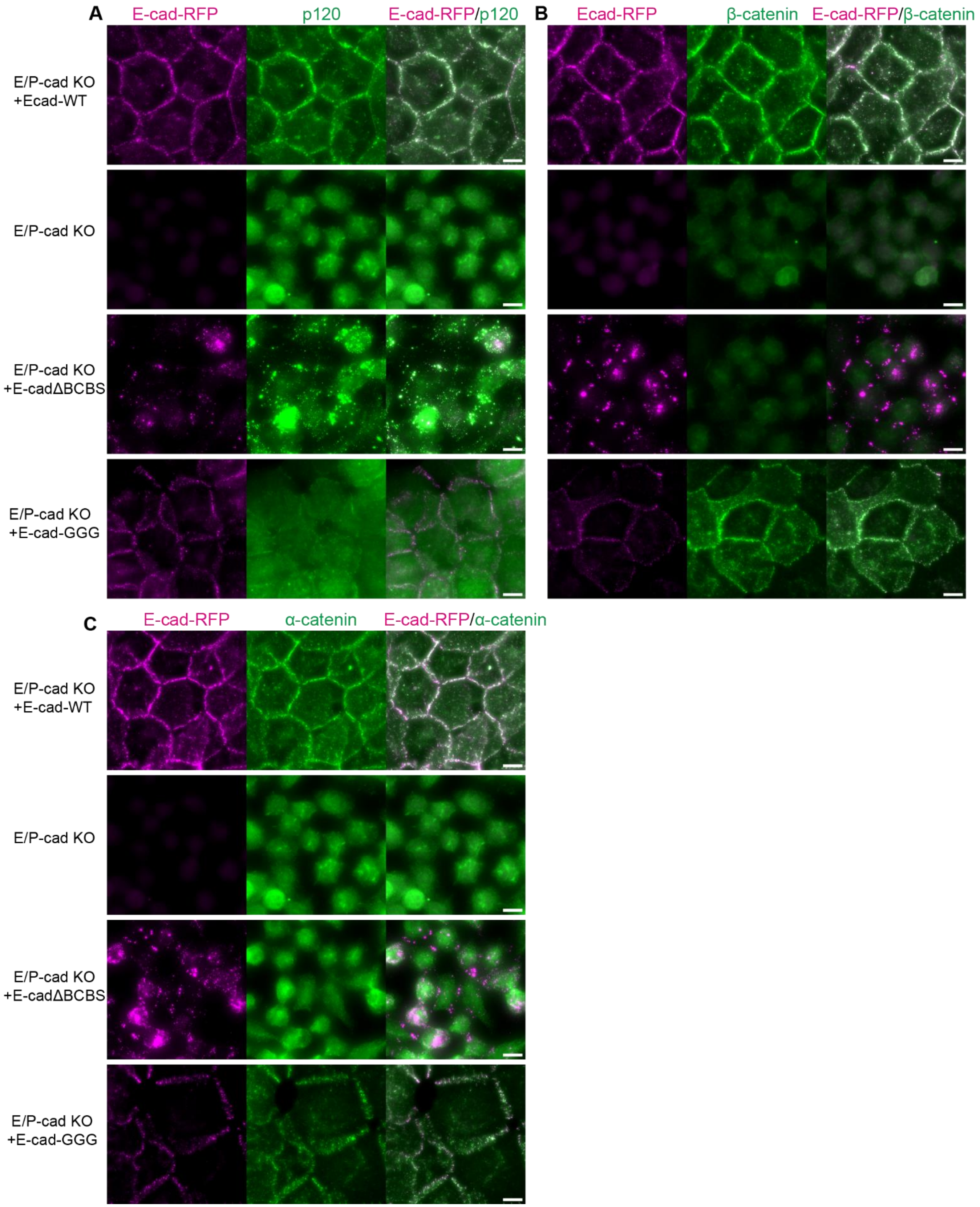


**Figure S1**: **Adherens junction protein localization in A431 E/P-cad KO cells expressing E-cadherin mutant proteins**. Fixed-cell immunofluorescence images of **A**) p120-catenin, **B**) β-catenin, and **C**) α-catenin in A431 E/P-cad KO cells expressing RFP-tagged E-cadherin mutants.

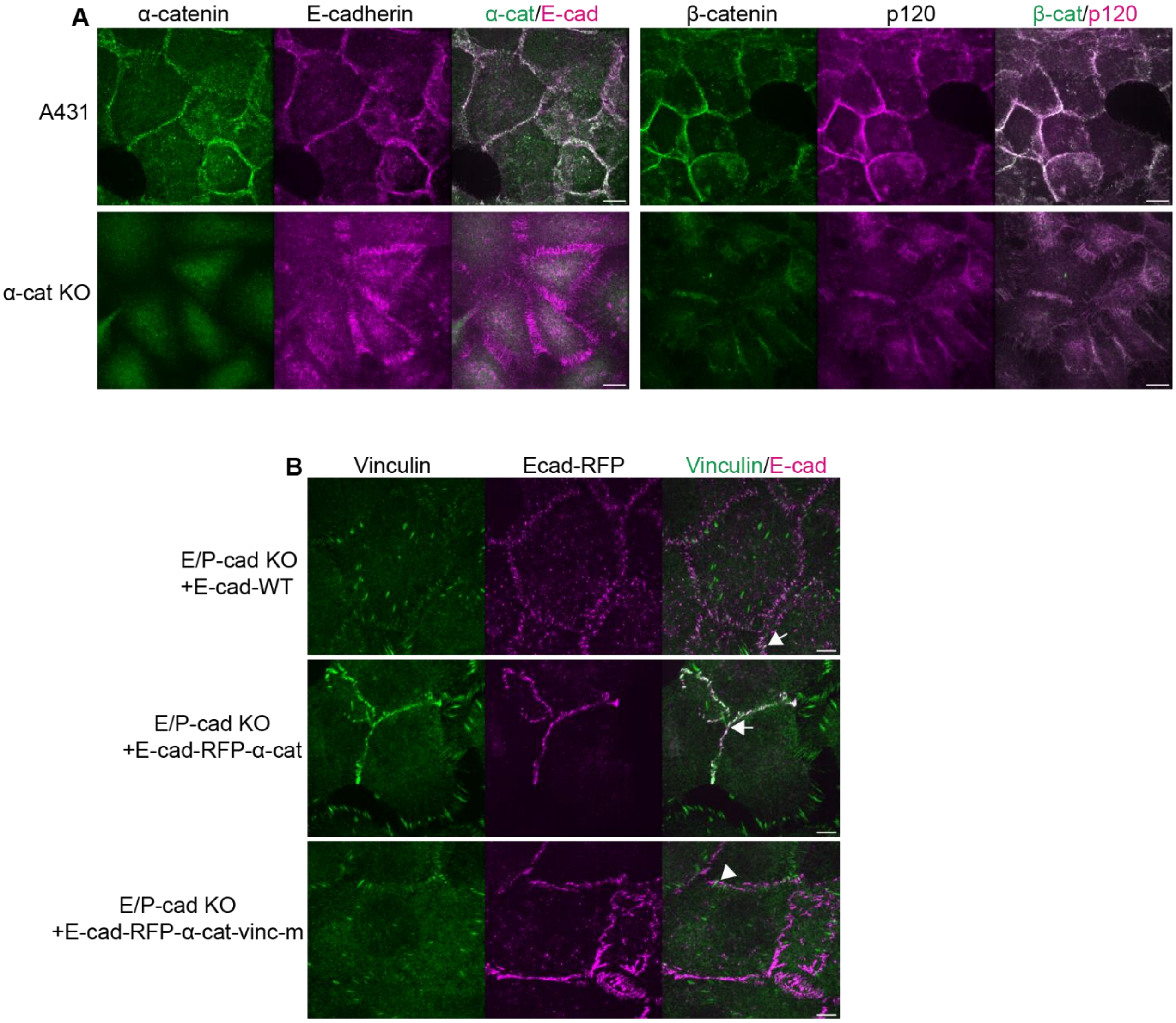


**Figure S2**: **Adherens junction protein localization in A431 α-catenin KO cells and E/P-cad KO cells expressing E-cadherin-α-catenin chimeric proteins. A)** Fixed-cell immunofluorescence images of α-catenin, E-cadherin, β-catenin, and p120-catenin in wild-type A431 and in A431 α-catenin KO cells. **B)** Fixed-cell immunofluorescence images of vinculin and E-cad-RFP localization in A431 E/P-cadherin knockout cells expressing different RFP-tagged E-cadherin mutants. In both E/P-cad KO+E-cad-WT and E/P-cad KO+Ecad-RFP-α-cat cells, vinculin colocalize with E-cad-RFP at adherens junctions (white arrows). However, in the E/P-cad KO+Ecad-RFP-α-cat-vinc-m cells— in which vinculin-binding domain in α-catenin is mutated—vinculin does not colocalize with E-cad-RFP at adherens junctions (white arrow).

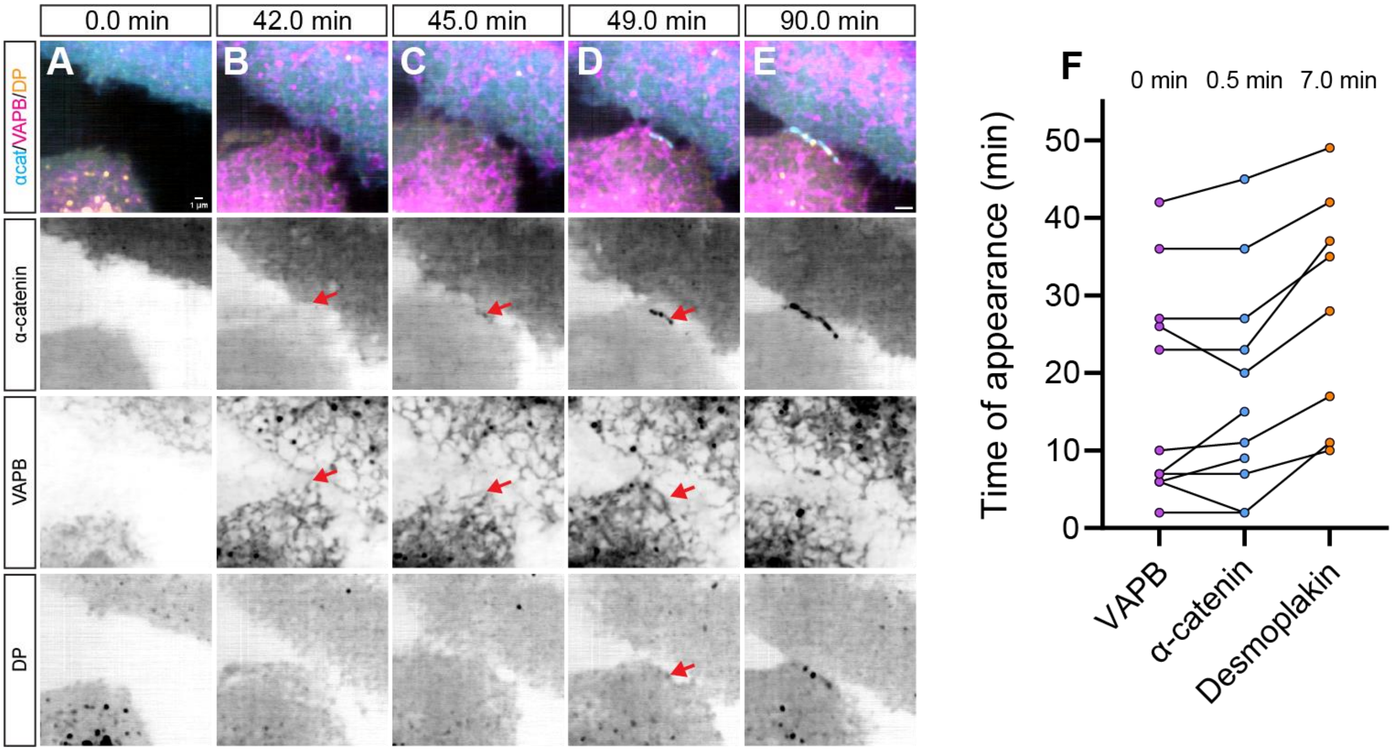


**Figure S3: Adherens junctions and ER appear simultaneously at nascent cell-cell interfaces, followed by desmosome assembly. A-E)** Live cell, time-lapse images of α-catenin, VAPB, and desmoplakin localization during cell-cell contact formation in A431 cells expressing mChilada-α-catenin, mApple-VAPB, and DP-EGFP. ER tubules are initially away from plasma membrane (**A**). α-catenin and VAPB appear within a few minutes of each other at newly formed cell-cell interface (**B-C, red arrows**). Desmoplakin appears after α-catenin and VAPB (**D**, **red arrows**). **F**) Time-course plot indicating the first appearance of VAPB, α-catenin, and desmoplakin at a calcium-induced nascent cell-cell contact formation. Each line corresponds to one sequence of cell-cell contact formation. The numbers above the graph indicate the mean time of the protein's first appearance. n=12 cell-cell contact formations.

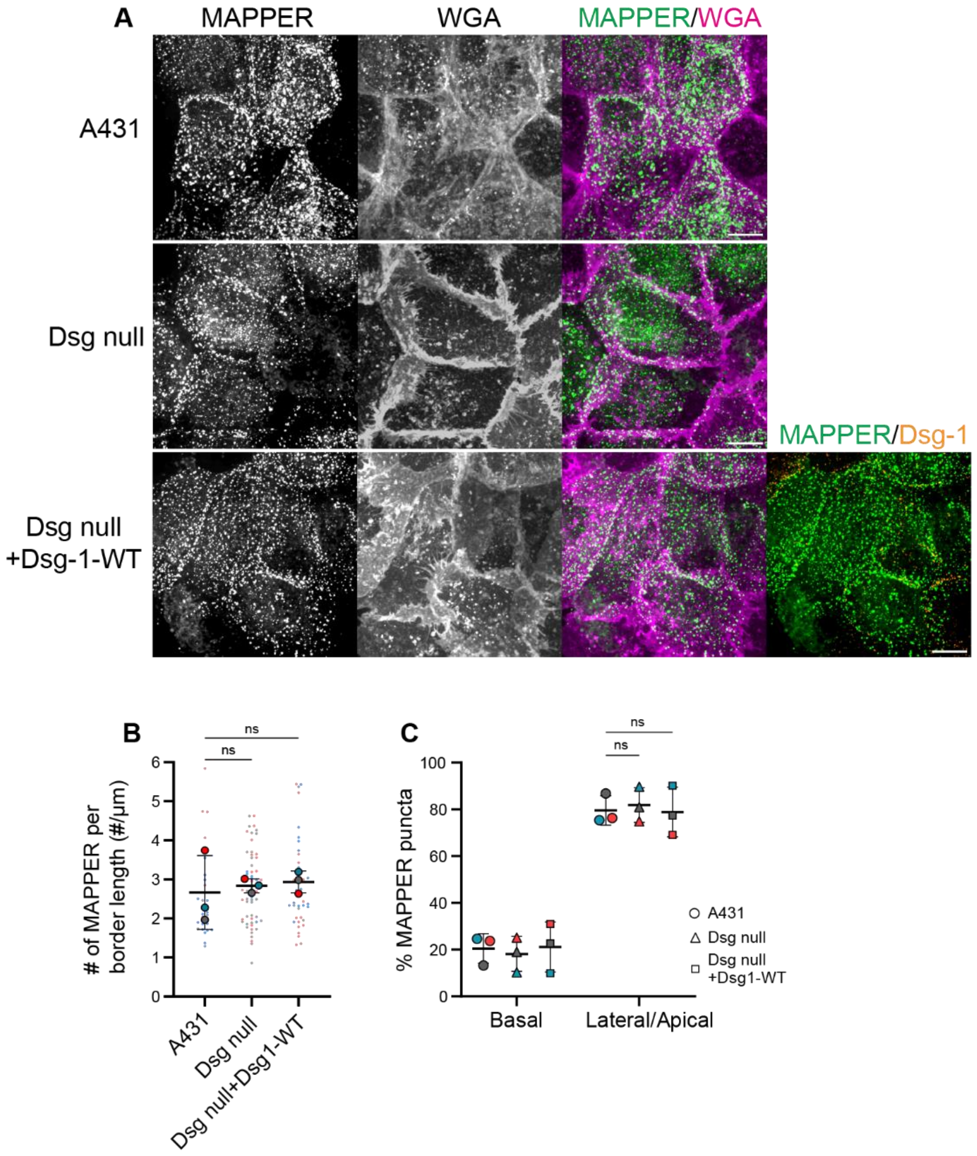


**Figure S4**: **Desmosome ablation does not impact ER-PMCS formation**. **A**) Live cell, maximum intensity projection images of ER-PM contact site marker mApple-MAPPER and membrane stain WGA-640R in wild-type A431, Dsg null, and Dsg null+Dsg-1-WT cells. Scale bar = 10 µm. **B**) The number of MAPPER puncta at cell-cell border, normalized to the length of border measured. color=experiment number; small dot=value for individual cell; large dot=mean value in each experiment; solid horizontal line=average of the 3 mean values; error bar=SD. p values=One-way ANOVA and Dunnett’s test. **C**) Percentage of total MAPPER puncta that is defined as basal or lateral/apical. color=experiment number; solid horizontal line=average of the 3 mean values; error bar=SD. p values=One-way ANOVA and Dunnett’s test.
